# A century of sampling at an ecological preserve reveals declining diversity of wild bees

**DOI:** 10.1101/2023.01.15.524123

**Authors:** Kelsey K. Graham, Paul Glaum, Joseph Hartert, Jason Gibbs, Erika Tucker, Rufus Isaacs, Fernanda S. Valdovinos

## Abstract

We analyzed the wild bee community from 1921 to 2018 at a nature preserve in southern Michigan, USA using museum records and found significant shifts in the bee community. Across the near century of records, species richness peaked in the 1970s and 1980s. There was an intensive bee survey completed by F.C. Evans in 1972 and 1973. We attempted to replicate his effort in 2017 and 2018, and again found a significant decline in species richness and evenness. There was also evidence of declining abundance in many of the more common species. We also conducted traits analyses using neural networks, revealing that oligolectic ground-nesting bees and cleptoparasitic bees were more likely to be extirpated whereas polylectic cavity-nesting bees were more likely to have persisted. Additionally, larger body size was associated with increased probability of local extirpation for polylectic cavity-nesting species. Larger phenological range was associated with increased chances of persistence for polylectic species, while it was associated with extirpation for oligolectic ground-nesting species. Species in the contemporary samples also had a more southerly overall distribution compared to the historic one.

**Open Research Statement:** Data used for analyses in this manuscript, including Evans’ original dataset from 1972/1973 with updated species nomenclature, will be permanently archived at the [USDA Ag Data Commons](https://data.nal.usda.gov/search/type/dataset) after the acceptance of this manuscript and will be citable and accessible here: https://data.nal.usda.gov/dataset/century-sampling-ecological-preserve-reveals-declining-diversity-wild-bees. Complete instructions on how to access all data referenced in this manuscript can be found in Appendix S1.

## Introduction

Pollinator health is a global concern, but our understanding of wild bee population declines is limited by a lack of long term monitoring (Woodard et al. 2020). Detecting population trends may require regular and intensive sampling over a long period (Williams et al. 2001). Consequently, many studies of bee population trends have analyzed datasets aggregated across multiple collection events to examine trends in the whole bee community (e.g. Bartomeus et al. 2013, Powney et al. 2019) or in specific groups of insects of interest such as bumble bees (Jacobson et al. 2018, Wood et al. 2019). Other alternatives to long-term monitoring include modeling of habitat suitability for wild bees (Koh et al. 2016) and comparing museum specimens from historical sampling over time (Bartomeus et al. 2018, Jacobson et al. 2018, Wood et al. 2019, Mathiasson and Rehan 2019).

Long-term monitoring programs in the United States are rare but, entomologists have sampled a few sites repeatedly due to interest in their fauna. These sites have been illuminating for understanding wild bee community changes. For example, in Illinois, Charles Robertson’s collection of bees and plants from natural habitats around Carlinville provided a foundation for contemporary sampling that enabled Burkle et al. (2013) to explore changes in plant-pollinator interaction networks over 120 years. Their study discovered a loss of bee species, temporal mismatches in plant-pollinator interactions, and the disruption of pollinator networks, predicting greater destabilization under future climate change scenarios.

The Edwin S. George Reserve (ESGR) is a 500 hectare preserve in southeast Michigan. It was originally cleared for agriculture before 1850 and cultivated until 1925–1926 (Evans 1986). Since then, it has not been managed by fire, cultivated for crops, or grazed by livestock. It was established as an ecological research station of the University of Michigan in 1930, with human activity limited to research activities and access road maintenance. From 1957 to 1989, bees and their floral associations at the ESGR were frequently sampled and recorded by Francis C. Evans, a faculty member at the University of Michigan, with particular intensity in 1972 and 1973 (Evans 1975, 1986). Evans’ sampling efforts spurred the National Research Council (2007) to highlight the ESGR as an appropriate site for addressing long-term trends in US pollinators.

Since Evans’ last efforts in 1989, only occasional records exist from 1990–1999, with no focused survey efforts. Prior to this study, there were no surveys or records for the ESGR after 1999.

We developed this study to: 1) determine whether contemporary resampling of the ESGR would reveal changes in the wild bee community or individual species historically at this site, and 2) evaluate if long-term bee community changes can be detected using the museum records available for the ESGR.

## Methods

Detailed methods are provided in Appendix S1. Briefly, an open area on the north side of the Edwin S. George Reserve in Livingston County, Michigan, USA (GPS coordinates: 42.456213, -84.018766) was the primary site for this study as it corresponds to the location of “Evans’ Old Field”, one of the areas historically sampled for bees. The site was visited every other week during the summers of 2017 and 2018. During each visit we 1) sampled bees along a 100 m transect for 40 minutes, 2) sampled bees randomly throughout the field (not restricted to the transect) for an additional 20 minutes, 3) spent 30 minutes sampling bees from each of the primary plants in bloom. All bees were identified to species (or lowest possible taxonomic level).

The University of Michigan Museum of Zoology Insect Collection (UMMZI), Ann Arbor, MI, holds over 4,000 bee specimens from historical collections at the ESGR. As part of this study, we worked with the UMMZI to catalog and transcribe label data from these specimens. Historical data were checked for entry errors and outdated taxonomies. Historical specimens whose taxon concept may have changed (e.g. *Lasioglossum*) were re-examined.

Specimens with questionable determinations were re-examined where possible, with difficult specimens loaned to the Gibbs Lab (University of Manitoba; current author) for identification.

In addition to analyses of the 4,000 plus records from the ESGR since 1921, we also conducted more focused analyses comparing Evans’ dataset from his 1972 and 1973 collection effort to our contemporary collections in 2017 and 2018. Evans’ original dataset was available through University of Michigan records, and we used these for our focused analyses comparing the contemporary sampling effort (2017–2018) to the historical one (1972–1973). The dataset is unique compared to the records from the museum, because Evans did not always collect observed bees if he was confident in their identification (Evans 1986). Therefore, his original dataset provides a more complete representation of the community he encountered.

## Data analyses

### Community change across a century of data

We employed an approach similar to that used by Bartomeus and colleagues (2013). To account for changes in practices across a century of records (1921–2018), such as including only a synoptic collection in the museum, we removed any duplicate specimen records from this data. These were records of the same species collected on the same date. Then, to account for uneven sampling, we binned specimen records into seven groupings of consecutive years to achieve more even sample numbers in each bin prior to comparison of species richness (Fig. S1). We then rarefied all bins to the bin with the lowest sample size (n = 320) using rarefy (package: vegan) to calculate the mean species richness and standard error of the mean. We used an analysis of variance (ANOVA) with a Tukey’s means difference test to determine significant differences between the bins.

### Comparison of contemporary and historical data sets

We compared the contemporary sampling dataset (years 2017 and 2018) to Evans’ original dataset from 1972 and 1973. We chose these years to compare due to similarities in sampling methods and intensity. We rarefied the species richness for the historical data and the contemporary data (rarefy, package: vegan) to the lowest sample size (contemporary data: n = 1271), and compared mean (+/- SE) rarefied species richness between the historical and the contemporary sampling years using an unpaired two-tailed t-test. We also calculated Shannon diversity (Hill number) for each sampling effort (hill_div; package: hilldiv) (Alberdi 2019), which accounts for both species richness and evenness in the population. Shannon diversity was statistically compared between datasets using Hutcheson t-test for two communities (Hutcheson_t_test, package: ecolTest) (Salinas 2021).

Additionally, we calculated Pielou’s evenness index for both datasets.

Non-metric multidimensional scaling (NMDS) ordinations were used to visualize differences in species composition between the historical and contemporary datasets (vegan package; Oksanen et al. 2019), and permutational multivariate analysis of variance (PERMANOVA) (adonis, package: vegan) (Anderson 2001) to determine whether the bee communities were significantly dissimilar in ordination between the two sampling periods. To compare dominant species within the two communities, we calculated the percent composition of each species to the overall community in the historical sampling years and the contemporary years. We then calculated the percent change in relative abundance for species with at least 30 records across the two sampling periods. Species were designated as declining if they had an abundance decline of more than 30% and increasing if they had an increase of more than 30%. Species in between were considered stable. This cutoff was chosen to match the IUCN designation for vulnerable species (IUCN Standards and Petitions Subcommittee 2014).

### Measuring change in community traits between contemporary and historical sampling periods

Species natural history traits were collected and collated for the species in our study (Table S1). We established latitudinal and longitudinal ranges per species and phenological ranges (flight periods) with specimen data available on the Global Biodiversity Information Facility (GBIF). We sourced intertegular distance values (ITD; mm) from published data (Gibbs et al. in press,

Castillo and Fairbairn 2012, Bartomeus et al. 2013, Cariveau et al. 2016, Lerman and Milam 2016, Hung 2017, Normandin et al. 2017, Nicholson 2019, Kendall et al. 2019, Lim et al. 2022) (Table S1). We used trait data in two types of analyses: 1) At the community level, we compared trait composition between the two periods’ sample communities, and 2) At the species level, we used traits as potential predictors of species persistence or local extirpation in the ESGR.

Trait composition differences between the historical and contemporary communities were initially explored via NMDS and PERMANOVA. However, the high dimensionality of the trait dataset limited further insight via traditional statistics. We therefore used neural networks due to their flexible non-parametric nature, ability to handle both categorical and continuous data, and increased capability to capture variance and interacting effects despite lingering collinearity and limited data. Detailed methods for our use of neural networks are provided in Appendix S1 (Fig S2:S3). Briefly, we used shallow (single layer) neural networks (nnet R package; Venables and Ripley 2002), as they maintain capability when data are limited for each class targeted for prediction (Basha et al. 2020). We estimated each trait’s (i.e., predictor’s) relative importance to model performance using the Garson method (NeuralNetTools package; Beck 2018). We then used the Olden method (NeuralNetTools package) to estimate the effect of each trait (predictor) on influencing specific model predictions. Finally, we tested/vetted sufficiently strong effects discovered via neural network analysis with linear frequentist statistics.

## Results

We found a unimodal species richness distribution over time, peaking in the 1970s (Table 1; see also Fig. S4). Rarefied bee species richness ranged from 87.8 ± 2.2 SE in Bin 1 (years: 1921-1959) to 110.6 ± 2.6 SE in Bin 6 (1982-1989). There was a significant difference in rarefied species richness among the bins (R^2^ = 0.021, F_6, 2757_ = 9.76, p < 0.001), peaking in bins 5 (years 1975-1981) and 6 (1982-1989) (Table 1). Select species abundance trends are in Fig S5.

**Table 1.** Rarefied species richness across binned years. Superscript letters indicate bins that are significantly different from each other (statistical results and reporting for the ANOVA with Tukey’s means difference analysis can be found in Table S2).

| Bin | Number of specimens | Years | Number of sampling days | Species Richness | Rarefied SR (mean +/- SE) |
| --- | --- | --- | --- | --- | --- |
| 1 | 385 | 1921-1959 | 174 | 94 | 87.8 ± 2.2<sup>a</sup> |
| 2 | 362 | 1960-1971 | 148 | 100 | 95.6 ± 1.9<sup>ab</sup> |
| 3 | 557 | 1972 | 61 | 122 | 103.0 ± 3.3<sup>bcd</sup> |
| 4 | 375 | 1973-1974 | 66 | 106 | 99.9 ± 2.1<sup>bc</sup> |
| 5 | 375 | 1975-1981 | 110 | 115 | 109.3 ± 2.1<sup>cd</sup> |
| 6 | 390 | 1982-1989 | 128 | 120 | 110.6 ± 2.6<sup>d</sup> |
| 7 | 320 | 1990-2018 | 40 | 98 | 98.0 ± 0.0<sup>ab</sup> |

Rarefied bee species richness was significantly greater in the historical sampling years (1972–1973) with an average of 119.1 ± 3.2 SE species compared to the contemporary sampling years (2017–2018) with an average of 91.0 ± 0.0 species (t = 6.22, df = 3835, p < 0.001). Shannon diversity (Hill’s number) was also significantly greater for the historical data (59 species) compared to the contemporary data (21 species) (*X*^2^ = 18.05, df = 1, p < 0.001). Both community estimates indicate that the wild bee population has fewer species and less population evenness in the contemporary community compared to the historical one.

There were 77 species collected in the historical sampling that were not collected in the contemporary sampling. There were also 33 species collected in the contemporary sampling effort that were not collected in the historical effort (Table S3). Additionally, the historical and contemporary bee communities were significantly different from each other in community composition (PERMANOVA; R^2^ = 0.26, F_1,9_ = 2.87, p = 0.01) (Fig. 1a). Seven species significantly (p < 0.01) contributed to ordination of the NMDS using envfit: *Lasioglossum pilosum, Lasioglossum lineatulum, Ceratina dupla, Halictus confusus, Halictus rubicundus, Colletes simulans*, and *Andrena hirticincta*. All of these species were less abundant in the contemporary sampling years compared to the historical sampling years (Fig. 1a).

**Figure 1.**
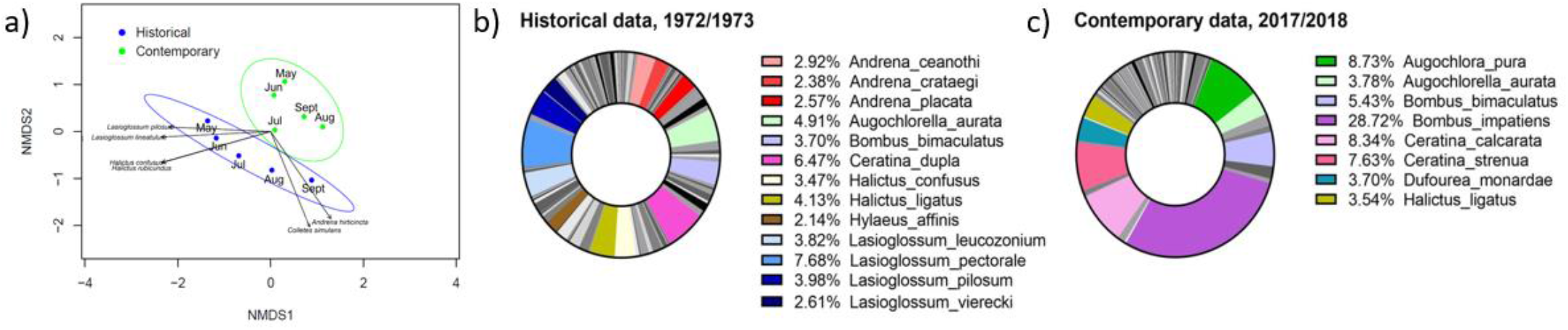
Community composition between periods. *a)* Visual representation of dissimilarity between the bee community trait composition during the historical (1972/1973) and the contemporary (2017/2018) bee collections, based on Bray-Curtis dissimilarity. Individual dots represent data collection months and ellipses indicate the 95% confidence intervals. Arrows indicate the most significant bee species driving the ordination of points. Arrow lengths correspond to the strength of the correlation between the species and the ordination. *b)* Proportions of species identified in the wild bee communities at the E.S. George Reserve during Evans’ data collection (historical: 1972/1973) and *c)* the contemporary samples (2017/2018). Species that represented more than 2% of the community are listed and colored in the chart.

The historical data had more species that represented over 2% of the specimens collected, as supported by a greater evenness index in the historical data (Pielou: 0.83) compared to the contemporary data (Pielou: 0.68). The most abundant species in the historical data were *Lasioglossum pectorale* (8% of specimens) and *C. dupla* (6% of specimens) (Fig. 1b). Decreased evenness in the contemporary data was especially driven by the increase in *Bombus impatiens*, which represented 29% of the contemporary specimens. *Augochlora pura* (9% of specimens), *Ceratina calcarata* (8%) and *Ceratina strenua* (8%) were also abundant (Fig. 1c).

There were 58 bee species collected in both the historical and contemporary sampling efforts. Only 22 of these had at least 30 records across the two sampling periods and were thus included in the calculation of relative abundance change (Table S4). Within this subset of species, 64% had a decline in relative abundance (>30% decrease), with an average percent decrease of 81% (± 5.1 S.E.). Fourteen percent of the species were considered stable, and 23% of species were increasing in relative abundance, with an average percent increase of 1130% (± 612.8 S.E.). Species with the greatest declines included *L. pilosum, H. confusus, C. dupla, Hylaeus affinis*, and *L. pectorale*, which all had a more than a 95% decline in relative abundance (Table S4). Species with the greatest increases included *A. pura, Lasioglossum versatum, B. impatiens*, and *Bombus citrinus*, which all increased by more than 1000% (Table S4).

At the species level, cavity nesting and polylectic species were most likely to persist compared to ground nesting and oligolectic species (Fig. 2a-d). Almost all oligolectic species were also ground nesting. Given sharp distinctions in persistence across our categorical traits, we used separate neural networks to study the relationship between quantitative traits and species persistence for each categorical subset in our data: 1) oligolectic ground nesting, 2) polylectic ground nesting, 3) polylectic cavity nesting, 4) cleptoparasitic (see Fig S6-S11 in SI Methods for details). This revealed that polylectic cavity-nesting and ground-nesting species were more likely to persist as phenological range increased (Fig. 2e, Figs. S4, S7). While oligolectic ground-nesters saw increased probability of extirpation with increasing phenological range (Fig. 2e, Fig. S5). Additionally, for polylectic cavity-nesters, bees were more likely to be extirpated with increasing ITD (Fig. S8). For the remaining groups and traits, results showed no effects, non-monotonic effects, or weak effects without verifiable statistical significance.

**Figure 2.**
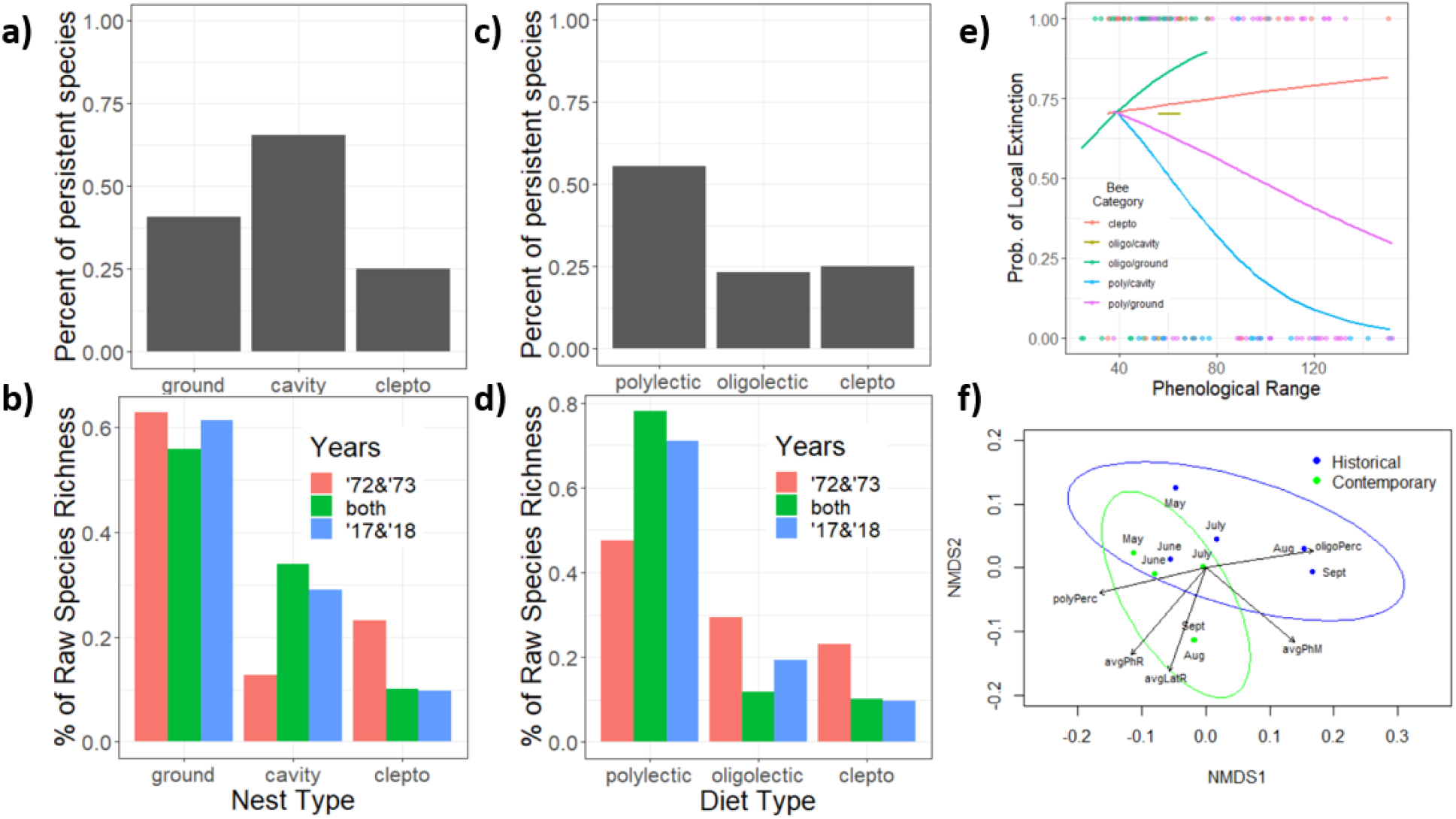
Trait changes between periods. *a & c)* Percent persistence of species separated by categorical traits across raw species counts. *b & d)* Percent of raw species richness across categorical traits grouped into species found only in the historical, in both, or only in the contemporary sampling effort. e) Visualized GLMM output showing the unique effects of Phenological Range (flight period length) and local extirpation probability across our different bee categories (see Table S7 for details). GLMM output mirrors neural net results, serving here to simply vet and visualize our conclusions. f) Visual representation of dissimilarity between the bee community trait composition during the historical (1972/1973) and the contemporary (2017/2018) bee collections, based on Bray-Curtis dissimilarity. Individual dots represent data collection months and ellipses indicate the 95% confidence intervals. Arrows indicate the most significant bee traits driving the ordination of points. Arrow lengths correspond to the strength of the correlation between the traits and the ordination.

At the community level, we observed clear differences between trait compositions in the historical (1972–1973) and contemporary (2017–2018) sampled communities (Fig. 2f, S12, S13). For instance, phenological range in the contemporary sample is significantly larger than the historical sample for all diet and nesting categories except oligolectic (Fig. S12). Additionally, the contemporary community had smaller average maximum latitude (∼3 degrees; Kruskal-Wallis *X*^2^=144.84, df = 1, p<2.2e-16) and minimum latitude (∼7 degrees; Kruskal-Wallis *X*^2^=320.10, df = 1, p< 2.2e-16), indicating higher relative abundances of bees with more southerly ranges in the 2010s than the 1970s.

## Discussion

We found evidence of significant bee community change over five decades, including a decline in species richness and evenness. The contemporary community was dominated by a few species, particularly *Bombus impatiens*. This is considered a species of least conservation concern, because it is increasing in abundance while also expanding its range (Cameron et al. 2011, Colla et al. 2012, Jacobson et al. 2018). For many other bee species, their abundance declined during the past few decades, with over half (64%) of analyzed species exhibiting a greater than 30% decline in relative abundance. Additionally, 58% of species recorded in the ESGR since 1921 were not captured in the 2017–2018 samples (Table S6). Phylogenetic groups did not appear to share similar fates, with congeners having vastly different trends in abundance.

Due to their ease of collection and prevalence in museum collections, *Bombus* species have been a focal group for measuring abundance and distribution trends (Grixti et al. 2009, Cameron et al. 2011, Jacobson et al. 2018, Wood et al. 2019). Similar to previous studies (Wood et al. 2019), we did not detect *B. affinis* in the contemporary sampling effort, and found that *B. bimaculatus* and *B. impatiens* are increasing in abundance, whereas *B. griseocollis* is stable.

Wood et al. (2019) found that *B. perplexus* and *B. vagans* are declining in distribution within the state, while at the ESGR we found they were stable. This may reflect the protected status of this site, with minimal human impacts since its establishment as an ecological reserve in 1930.

Other species with significant declines in relative abundance include two *Lasioglossum* species (*L. pilosum* and *L. pectorale*), *Halictus confusus, Ceratina dupla* and *Hylaeus affinis*. In another study of species trends in west Michigan, *L. pilosum, L. pectorale, and H. confusus* were stable in abundance for over sixteen years (Graham et al. 2021). All five of these species are widespread and abundant in numerous biodiversity surveys in eastern North America (e.g. Mallinger et al. 2016, Choate et al. 2018). This highlights how location-specific species population trends can be. Though evidence of species declines across diverse locations and landscapes may indicate more widespread threats to species, including threats of climate change. There is some evidence of climate change being a driver of community composition, as bees in the 2017-2018 samples had more southerly ranges.

Ecological or life history traits have been used previously to understand what makes a species vulnerable. Regarding lecty, our data are similar to past studies (Bartomeus et al. 2013, Burkle et al. 2013), with oligolectic bees being the most likely to be extirpated, while polylectic cavity-nesting bees were more likely to persist. Burkle and colleagues (2013) found a greater loss of cavity-nesting species, whereas we found they were more likely to persist. However, their study occurred in a predominately grassland dominated landscape heavily influenced by agricultural intensification, while our study was in a forested ecological preserve. For polylectic cavity-nesting species, we found that increasing ITD (larger bodied bees) were more strongly associated with extirpation and bees with large phenological ranges were associated with persistence, similar to Bartomeus et al. (2013). We also found that for oligolectic ground-nesting bees, larger phenological ranges were associated with extirpation. Oligolectic ground-nesting bees were the group with the highest proportion of extirpated species between 1972–1973 and 2017–2018 (78%), which could be driven by the encroachment of autumn olive (*Elaeagnus umbellata*) at the preserve and decreasing overall open space where sampling was focused.

This study provides strong evidence of recent wild bee community change at a nature preserve since the early 1970s, with a decline in species richness, evenness, and abundance. Traits analyses identified groups that are more likely to be locally extirpated, i.e., oligolectic ground-nesting and cleptoparastic bees, and those that are more likely to persist, i.e., polylectic cavity-nesting species. We also found evidence of range shifts, with bees in more southerly ranges increasing in abundance.

## Acknowledgements

We thank Dr. Christopher Dick, Robyn Burnham and Alex Wenner for access to the ESGR. We thank Joel Gardner (University of Manitoba) for his assistance identifying bees. We also thank the many undergraduates and technicians that helped with this research, particularly Michael Killewald, Alexandrea Peake, Katie Manning, Ryan Oleynik, Peregrine Ke-Lind, Grace Haynes, and Meagan Sevener. This project was funded by US Department of Agriculture grant number 2017-68004-26323 from the National Institute of Food and Agriculture (R. Isaacs, J. Gibbs) and National Science Foundation grants DEB-1834497 and DEB-2129757 (F.S. Valdovinos).

## References

Alberdi, A. 2019. Integral Analysis of Diversity Based on Hill Numbers.

Anderson, M. J. 2001. A new method for non-parametric multivariate analysis of variance. Austral Ecology 26:32–46.

Bartomeus, I., J. S. Ascher, J. Gibbs, B. N. Danforth, D. L. Wagner, S. M. Hedtke, and R. Winfree. 2013. Historical changes in northeastern US bee pollinators related to shared ecological traits. Proceedings of the National Academy of Sciences of the United States of America 110:4656–60.

Bartomeus, I., J. R. Stavert, D. Ward, and O. Aguado. 2018. Historical collections as a tool for assessing the global pollination crisis. Philosophical Transactions of the Royal Society B 374.

Beck, M. W. 2018. NeuralNetTools: Visualization and Analysis Tools for Neural Networks. Journal of Statistical Software 85:1–20.

Burkle, L. A., J. C. Marlin, and T. M. Knight. 2013. Plant-pollinator interactions over 120 years: loss of species, co-occurrence, and function. Science 339:1611–1615.

Cameron, S. A., J. D. Lozier, J. P. Strange, J. B. Koch, N. Cordes, L. F. Solter, and T. L. Griswold. 2011. Patterns of widespread decline in North American bumble bees. Proceedings of the National Academy of Sciences of the United States of America 108:662–7.

Cariveau, D. P., G. K. Nayak, I. Bartomeus, J. Zientek, J. S. Ascher, J. Gibbs, and R. Winfree. 2016. The Allometry of Bee Proboscis Length and Its Uses in Ecology. PLOS ONE 11:e0151482.

Castillo, R. C. del, and D. J. Fairbairn. 2012. Macroevolutionary patterns of bumblebee body size: detecting the interplay between natural and sexual selection. Ecology and Evolution 2:46–57.

Choate, B. A., P. L. Hickman, and E. A. Moretti. 2018. Wild bee species abundance and richness across an urban–rural gradient. Journal of Insect Conservation 22:391–403.

Colla, S. R., F. Gadallah, L. Richardson, D. Wagner, and L. Gall. 2012. Assessing declines of North American bumble bees (Bombus spp.) using museum specimens. Biodiversity and Conservation 21:3585–3595.

Committee on the Status of Pollinators in North America - National Research Council. 2007. Status of Pollinators in North America.

Evans, F. C. 1975. The natural history of a Michigan field. Pages 27–51 in M. K. Wali, editor. Prairie: a Multiple View. University of North Dakota Press, Grand Forks, ND.

Evans, F. C. 1986. Bee - Flower Interactions on an Old Field in Southeastern Michigan. Pages 103–109 in G. K. Clambey and R. H. Pembley, editors. The prairie: past, present and future. Proceedings of the ninth North American Prairie Conference. Tri-College University Center for Environmental Studies, Fargo, ND.

Gibbs, J., E. Hanuschuk, R. Miller, M. Dubois, M. Martini, S. Robinson, P. Nakagawa, C. Sheffield, S. Cardinal, and T. Onuferko. (in press). A checklist of the bees (Hymenoptera: Apoidea) of Manitoba. The Canadian Entomologist.

Graham, K. K., J. Gibbs, J. Wilson, E. May, and R. Isaacs. 2021. Resampling of wild bees across fifteen years reveals variable species declines and recoveries after extreme weather. Agriculture, Ecosystems & Environment 317:107470.

Grixti, J. C., L. T. Wong, S. A. Cameron, and C. Favret. 2009. Decline of bumble bees (Bombus) in the North American Midwest. Biological Conservation 142:75–84.

Hung, K.-L. J. 2017. Effects of Habitat Fragmentation and Introduced Species on the Structure and Function of Plant-Pollinator Interactions. University of California, San Diego, San Diego, CA.

IUCN Standards and Petitions Subcommittee. 2014. Guidelines for Using the IUCN Red List Categories and Criteria. IUCN.

Jacobson, M. M., E. M. Tucker, M. E. Mathiasson, and S. M. Rehan. 2018. Decline of bumble bees in northeastern North America, with special focus on Bombus terricola. Biological Conservation 217:437–445.

Kendall, L. K., R. Rader, V. Gagic, D. P. Cariveau, M. Albrecht, K. C. R. Baldock, B. M. Freitas, M. Hall, A. Holzschuh, F. P. Molina, J. M. Morten, J. S. Pereira, Z. M. Portman, S. P. M. Roberts, J. Rodriguez, L. Russo, L. Sutter, N. J. Vereecken, and I. Bartomeus. 2019. Pollinator size and its consequences: Robust estimates of body size in pollinating insects. Ecology and Evolution 9:1702–1714.

Koh, I., E. V. Lonsdorf, N. M. Williams, C. Brittain, R. Isaacs, J. Gibbs, and T. H. Ricketts. 2016. Modeling the status, trends, and impacts of wild bee abundance in the United States. Proceedings of the National Academy of Sciences 113:140–145.

Lerman, S. B., and J. Milam. 2016. Bee Fauna and Floral Abundance Within Lawn-Dominated Suburban Yards in Springfield, MA. Annals of the Entomological Society of America 109:713–723.

Lim, K., S. Lee, M. Orr, and S. Lee. 2022. Harrison’s rule corroborated for the body size of cleptoparasitic cuckoo bees (Hymenoptera: Apidae: Nomadinae) and their hosts. Scientific Reports 2022 12:1 12:1–12.

Mallinger, R. E., J. Gibbs, and C. Gratton. 2016. Diverse landscapes have a higher abundance and species richness of spring wild bees by providing complementary floral resources over bees’ foraging periods. Landscape Ecology 31:1523–1535.

Mathiasson, M. E., and S. M. Rehan. 2019. Status changes in the wild bees of north-eastern North America over 125 years revealed through museum specimens. Insect Conservation and Diversity 12:278–288.

Nicholson, C. 2019. Nicholson_etal_2019_Wild_bee_occurrence_traits. figshare.

Normandin, É., N. J. Vereecken, C. M. Buddle, and V. Fournier. 2017. Taxonomic and functional trait diversity of wild bees in different urban settings. PeerJ 2017:e3051.

Oksanen, J., F. G. Blanchet, M. Friendly, R. Kindt, P. Legendre, D. McGlinn, P. R. Minchin, R. B. O’Hara, G. L. Simpson, P. Solymos, M. H. H. Stevens, E. Szoecs, and H. Wagner. 2019. vegan: Community Ecology Package.

Powney, G. D., C. Carvell, M. Edwards, R. K. A. Morris, H. E. Roy, B. A. Woodcock, and N. J. B. Isaac. 2019. Widespread losses of pollinating insects in Britain. Nature Communications 10:1018.

Salinas, H. 2021. ecolTest: Community Ecology Tests.

Venables, W. N., and B. D. Ripley. 2002. Modern Applied Statistics with S. Fourth. Springer, New York.

Williams, N. M., R. L. Minckley, and F. A. Silveira. 2001. Variation in Native Bee Faunas and its Implications for Detecting Community Changes. Conservation Ecology 5.

Wood, T. J., J. Gibbs, K. K. Graham, and R. Isaacs. 2019. Narrow pollen diets are associated with declining Midwestern bumble bee species. Ecology 100:e02697.

Woodard, S. H., S. Federman, R. R. James, B. N. Danforth, T. L. Griswold, D. Inouye, Q. S. McFrederick, L. Morandin, D. L. Paul, E. Sellers, J. P. Strange, M. Vaughan, N. M. Williams, M. G. Branstetter, C. T. Burns, J. Cane, A. B. Cariveau, D. P. Cariveau, A. Childers, C. Childers, D. L. Cox-Foster, E. C. Evans, K. K. Graham, K. Hackett, K. T. Huntzinger, R. E. Irwin, S. Jha, S. Lawson, C. Liang, M. M. López-Uribe, A. Melathopoulos, H. M. C. Moylett, C. R. V. Otto, L. C. Ponisio, L. L. Richardson, R. Rose, R. Singh, and W. Wehling. 2020. Towards a U.S. national program for monitoring native bees. Biological Conservation 252:108821.

