## Supplemental Information for "A century of sampling at an ecological preserve reveals declining diversity of wild bees"

### Supplementary Tables

**Table S1.** Traits data
Trait data is available on Data Dryad:
https://datadryad.org/stash/share/TVZGtBbRd3x1lt9x_VM5ZzRfBIEO1JgkKBWyzRsb38

**Table S2.** Results of Tukey's multiple comparisons test to compare bee species richness between binned years of specimens across a near century of sampling (1921-2018).

| | |
|---|---|
| **Number of families** | 1 |
| **Number of comparisons per family** | 21 |
| **Alpha** | 0.05 |

| Tukey's multiple comparisons test | Mean Diff. | 95.00% CI of diff. | Below threshold? | Summary | Adjusted P Value |
|---|---|---|---|---|---|
| Bin 1: 1921-1959 vs. Bin 2: 1960-1971 | -7.800 | -18.55 to 2.945 | No | ns | 0.3283 |
| Bin 1: 1921-1959 vs. Bin 3: 1972 | -15.20 | -24.93 to -5.472 | Yes | **** | <0.0001 |
| Bin 1: 1921-1959 vs. Bin 4: 1973-1974 | -12.10 | -22.75 to -1.451 | Yes | * | 0.0143 |
| Bin 1: 1921-1959 vs. Bin 5: 1975-1981 | -21.50 | -32.15 to -10.85 | Yes | **** | <0.0001 |
| Bin 1: 1921-1959 vs. Bin 6: 1982-1989 | -22.80 | -33.34 to -12.26 | Yes | **** | <0.0001 |
| Bin 1: 1921-1959 vs. Bin 7: 1990-2018 | -10.20 | -21.30 to 0.9029 | No | ns | 0.0959 |
| Bin 2: 1960-1971 vs. Bin 3: 1972 | -7.400 | -17.31 to 2.509 | No | ns | 0.2938 |
| Bin 2: 1960-1971 vs. Bin 4: 1973-1974 | -4.300 | -15.11 to 6.515 | No | ns | 0.9043 |
| Bin 2: 1960-1971 vs. Bin 5: 1975-1981 | -13.70 | -24.51 to -2.885 | Yes | ** | 0.0036 |
| Bin 2: 1960-1971 vs. Bin 6: 1982-1989 | -15.00 | -25.71 to -4.288 | Yes | *** | 0.0007 |
| Bin 2: 1960-1971 vs. Bin 7: 1990-2018 | -2.400 | -13.66 to 8.862 | No | ns | 0.9959 |
| Bin 3: 1972 vs. Bin 4: 1973-1974 | 3.100 | -6.704 to 12.90 | No | ns | 0.9673 |
| Bin 3: 1972 vs. Bin 5: 1975-1981 | -6.300 | -16.10 to 3.504 | No | ns | 0.4831 |
| Bin 3: 1972 vs. Bin 6: 1982-1989 | -7.600 | -17.29 to 2.091 | No | ns | 0.2375 |
| Bin 3: 1972 vs. Bin 7: 1990-2018 | 5.000 | -5.295 to 15.30 | No | ns | 0.7840 |
| Bin 4: 1973-1974 vs. Bin 5: 1975-1981 | -9.400 | -20.12 to 1.319 | No | ns | 0.1301 |
| Bin 4: 1973-1974 vs. Bin 6: 1982-1989 | -10.70 | -21.32 to -0.08476 | Yes | * | 0.0467 |
| Bin 4: 1973-1974 vs. Bin 7: 1990-2018 | 1.900 | -9.270 to 13.07 | No | ns | 0.9988 |
| Bin 5: 1975-1981 vs. Bin 6: 1982-1989 | -1.300 | -11.92 to 9.315 | No | ns | 0.9998 |
| Bin 5: 1975-1981 vs. Bin 7: 1990-2018 | 11.30 | 0.1301 to 22.47 | Yes | * | 0.0453 |
| Bin 6: 1982-1989 vs. Bin 7: 1990-2018 | 12.60 | 1.529 to 23.67 | Yes | * | 0.0140 |

| Test details | Mean 1 | Mean 2 | Mean Diff. | SE of diff. | n1 | n2 | q | DF |
|---|---|---|---|---|---|---|---|---|
| Bin 1: 1921-1959 vs. Bin 2: 1960-1971 | 87.80 | 95.60 | -7.800 | 3.642 | 385 | 362 | 3.029 | 2757 |
| Bin 1: 1921-1959 vs. Bin 3: 1972 | 87.80 | 103.0 | -15.20 | 3.297 | 385 | 557 | 6.520 | 2757 |
| Bin 1: 1921-1959 vs. Bin 4: 1973-1974 | 87.80 | 99.90 | -12.10 | 3.609 | 385 | 375 | 4.741 | 2757 |
| Bin 1: 1921-1959 vs. Bin 5: 1975-1981 | 87.80 | 109.3 | -21.50 | 3.609 | 385 | 375 | 8.424 | 2757 |
| Bin 1: 1921-1959 vs. Bin 6: 1982-1989 | 87.80 | 110.6 | -22.80 | 3.574 | 385 | 390 | 9.022 | 2757 |

| | | | | | | | | |
|---|---|---|---|---|---|---|---|---|
| Bin 1: 1921-1959 vs. Bin 7: 1990-2018 | 87.80 | 98.00 | -10.20 | 3.763 | 385 | 320 | 3.833 | 2757 |
| Bin 2: 1960-1971 vs. Bin 3: 1972 | 95.60 | 103.0 | -7.400 | 3.358 | 362 | 557 | 3.116 | 2757 |
| Bin 2: 1960-1971 vs. Bin 4: 1973-1974 | 95.60 | 99.90 | -4.300 | 3.665 | 362 | 375 | 1.659 | 2757 |
| Bin 2: 1960-1971 vs. Bin 5: 1975-1981 | 95.60 | 109.3 | -13.70 | 3.665 | 362 | 375 | 5.286 | 2757 |
| Bin 2: 1960-1971 vs. Bin 6: 1982-1989 | 95.60 | 110.6 | -15.00 | 3.631 | 362 | 390 | 5.843 | 2757 |
| Bin 2: 1960-1971 vs. Bin 7: 1990-2018 | 95.60 | 98.00 | -2.400 | 3.817 | 362 | 320 | 0.8892 | 2757 |
| Bin 3: 1972 vs. Bin 4: 1973-1974 | 103.0 | 99.90 | 3.100 | 3.323 | 557 | 375 | 1.319 | 2757 |
| Bin 3: 1972 vs. Bin 5: 1975-1981 | 103.0 | 109.3 | -6.300 | 3.323 | 557 | 375 | 2.681 | 2757 |
| Bin 3: 1972 vs. Bin 6: 1982-1989 | 103.0 | 110.6 | -7.600 | 3.284 | 557 | 390 | 3.272 | 2757 |
| Bin 3: 1972 vs. Bin 7: 1990-2018 | 103.0 | 98.00 | 5.000 | 3.489 | 557 | 320 | 2.026 | 2757 |
| Bin 4: 1973-1974 vs. Bin 5: 1975-1981 | 99.90 | 109.3 | -9.400 | 3.633 | 375 | 375 | 3.659 | 2757 |
| Bin 4: 1973-1974 vs. Bin 6: 1982-1989 | 99.90 | 110.6 | -10.70 | 3.598 | 375 | 390 | 4.206 | 2757 |
| Bin 4: 1973-1974 vs. Bin 7: 1990-2018 | 99.90 | 98.00 | 1.900 | 3.786 | 375 | 320 | 0.7098 | 2757 |
| Bin 5: 1975-1981 vs. Bin 6: 1982-1989 | 109.3 | 110.6 | -1.300 | 3.598 | 375 | 390 | 0.5110 | 2757 |
| Bin 5: 1975-1981 vs. Bin 7: 1990-2018 | 109.3 | 98.00 | 11.30 | 3.786 | 375 | 320 | 4.221 | 2757 |

**Table S3**. Raw abundances of bee species collected at the E.S George Reserve, Evans' Old Field in 1972-73 (historical) and in 2017-18 (contemporary). Red shading indicates species that were not detected in the contemporary sampling years, and yellow shading indicates species that were only found in the contemporary years, and not found in the historical collections.

| **Family** | **Species** | **Historical** | **Contemporary** |
|---|---|---|---|
| Andrenidae | *Andrena alleghaniensis* Viereck, 1897 | 4 | |
| | *Andrena barbilabris* (Kirby, 1802) | | 2 |
| | *Andrena brevipalpis* Cockerell, 1930 | | 1 |
| | *Andrena canadensis* Dalla Torre, 1896 | 11 | |
| | *Andrena carlini* Cockerell, 1901 | 1 | 1 |
| | *Andrena ceanothi* Viereck, 1917 | 75 | |
| | *Andrena commoda* Smith, 1879 | 3 | |
| | *Andrena crataegi* Robertson, 1893 | 61 | 3 |
| | *Andrena cressonii* Robertson, 1891 | 2 | |
| | *Andrena dunningi* Cockerell, 1898 | | 1 |
| | *Andrena forbesii* Robertson, 1891 | 10 | 9 |
| | *Andrena fragilis* Smith, 1853 | 3 | |
| | *Andrena hirticincta* Provancher, 1888 | 19 | 7 |
| | *Andrena imitatrix* Cresson, 1872 | 11 | |

| | | |
|---|---|---|
| *Andrena krigiana* Robertson, 1901 | | 14 |
| *Andrena mandibularis* Robertson, 1892 | 1 | |
| *Andrena melanochroa* Cockerell, 1989 | 2 | |
| *Andrena miranda* Smith, 1879 | 6 | |
| *Andrena miserabilis** Cresson, 1872 | | 6 |
| *Andrena nasonii* Robertson, 1895 | 4 | |
| *Andrena nivalis* Smith, 1853 | 1 | 1 |
| *Andrena nubecula* Smith, 1853 | 12 | 19 |
| *Andrena perplexa** Smith, 1853 | | 1 |
| *Andrena placata* Mitchell, 1960 | 66 | |
| *Andrena platyparia* Robertson, 1895 | 1 | 1 |
| *Andrena rudbeckiae* Robertson, 1891 | 10 | |
| *Andrena rugosa* Cockerell, 1891 | 3 | 3 |
| *Andrena salictaria* Robertson, 1905 | 1 | |
| *Andrena sigmundi* Cockerell, 1902 | 42 | |
| *Andrena vicina* Smith, 1853 | 32 | 6 |
| *Andrena wilkella* (Kirby, 1802) | 14 | |
| *Calliopsis andreniformis* Smith, 1853 | 25 | |

| | | | |
|---|---|---|---|
| | *Perdita bequaerti* Viereck, 1917 | 1 | |
| | *Perdita octomaculata* (Say, 1824) | 2 | |
| | *Pseudopanurgus aestivalis* Crawford, 1903 | 17 | |
| | *Pseudopanurgus andrenoides* Smith, 1853 | | 5 |
| Apidae | *Agapostemon sericeus* (Forster, 1771) | 3 | |
| | *Agapostemon splendens* (Lepeletier, 1841) | 8 | |
| | *Agapostemon texanus* (Cresson, 1872) | 19 | 1 |
| | *Agapostemon virescens* (Fabricius, 1775) | 24 | |
| | *Anthophora terminalis* Cresson, 1869 | 3 | 1 |
| | *Bombus affinis* Cresson, 1863 | 17 | |
| | *Bombus ashtoni* (Cresson, 1864) | 13 | |
| | *Bombus auricomus* (Robertson, 1903) | 18 | |
| | *Bombus bimaculatus* Cresson, 1863 | 95 | 69 |
| | *Bombus borealis* Kirby, 1837 | 4 | |
| | *Bombus citrinus* (Smith, 1854) | 3 | 25 |
| | *Bombus fernaldae* (Franklin, 1911) | 3 | |
| | *Bombus fervidus* (Fabricius, 1798) | 12 | |

| | | |
|---|---|---|
| *Bombus griseocollis* (DeGeer, 1773) | 19 | 9 |
| *Bombus impatiens* Cresson, 1863 | 29 | 365 |
| *Bombus perplexus* Cresson, 1863 | 8 | 4 |
| *Bombus terricola* Kirby, 1837 | 3 | |
| *Bombus vagans* Smith, 1854 | 33 | 16 |
| *Ceratina calcarata** Robertson, 1900 | | 106 |
| *Ceratina dupla* Say, 1837 | 166 | 2 |
| *Ceratina mikmaqi* Rehan & Sheffield, 2011 | | 11 |
| *Ceratina strenua* Smith, 1879 | | 97 |
| *Epeolus pusillus* Cresson, 1864 | 6 | |
| *Epeolus scutellaris* Say, 1824 | 15 | 3 |
| *Holcopasites calliopsidis* (Linsley, 1943) | 7 | |
| *Melissodes bimaculatus* (Lepeletier, 1825) | | 1 |
| *Melissodes communis* Cresson, 1878 | 6 | |
| *Melissodes desponsus* Smith, 1854 | 18 | 2 |
| *Melissodes druriellus* (Kirby, 1802) | 6 | |
| *Melissodes illatus** Lovell and Cockerell, 1906 | | 2 |
| *Melissodes niveus* Robertson, 1895 | 1 | |

| | | | |
|---|---|---|---|
| | *Melissodes subillatus* LaBerge, 1961 | 6 | |
| | *Melissodes tinctus* LaBerge, 1961 | 4 | 1 |
| | *Melissodes wheeleri* Cockerell, 1906 | 1 | |
| | *Nomada armatella* Cockerell, 1903 | 1 | |
| | *Nomada cressonii* Robertson, 1893 | 25 | 4 |
| | *Nomada cuneata* (Robertson, 1903) | | 1 |
| | *Nomada luteoloides* Robertson, 1895 | 1 | |
| | *Nomada maculata* Cresson, 1863 | 7 | 3 |
| | *Nomada pygmaea* Cresson, 1863 | 18 | |
| | *Nomada sayi* Robertson, 1893 | 1 | |
| | *Nomada subrutila* Lovell and Cockerell, 1905 | 2 | |
| | *Nomada vicina* Cresson, 1863 | 4 | |
| | *Peponapis pruinosa* (Say, 1837) | | 1 |
| | *Triepeolus donatus* (Smith, 1854) | 2 | |
| | *Xylocopa virginica* (Linnaeus, 1771) | 2 | 1 |
| Colletidae | *Colletes americanus* Cresson, 1868 | 19 | |
| | *Colletes compactus* Cresson, 1868 | 2 | |
| | *Colletes inaequalis* Say, 1837 | 4 | |

| | | | |
|---|---|---|---|
| | *Colletes latitarsis* Robertson, 1891 | 1 | |
| | *Colletes nudus* Robertson, 1898 | 2 | |
| | *Colletes simulans* Cresson, 1868 | 37 | 2 |
| | *Colletes solidaginis* Swenk, 1906 | 19 | |
| | *Colletes validus* Cresson, 1868 | 2 | |
| | *Hylaeus affinis* (Smith, 1853) | 55 | 1 |
| | *Hylaeus illinoisensis* (Robertson, 1896) | | 2 |
| | *Hylaeus mesillae* (Cockerell, 1896) | 28 | 1 |
| | *Hylaeus modestus* Say, 1837 | 9 | 2 |
| Halictidae | *Augochlora pura* (Say, 1837) | 7 | 111 |
| | *Augochlorella aurata* (Smith, 1853) | 126 | 48 |
| | *Augochlorella persimilis* (Viereck, 1910) | | 24 |
| | *Augochloropsis metallica* (Smith, 1853) | 37 | 13 |
| | *Dufourea monardae* (Viereck, 1924) | 31 | 47 |
| | *Halictus confusus* Smith, 1853 | 89 | 1 |
| | *Halictus ligatus* Say, 1837 | 106 | 45 |
| | *Halictus parallelus* Say, 1837 | 6 | |
| | *Halictus rubicundus* (Christ, 1791) | 42 | |

| | | |
|---|---|---|
| *Lasioglossum abanci* (Crawford, 1932) | | 2 |
| *Lasioglossum anomalum* (Robertson, 1935) | 9 | 6 |
| *Lasioglossum bruneri* (Crawford, 1902) | 1 | |
| *Lasioglossum cattellae* (Ellis, 1913) | | 15 |
| *Lasioglossum coeruleum** (Robertson, 1893) | | 1 |
| *Lasioglossum coriaceum* (Smith, 1853) | 42 | 7 |
| *Lasioglossum cressonii* (Robertson, 1890) | 13 | 11 |
| *Lasioglossum foveolatum* (Robertson, 1902) | 1 | |
| *Lasioglossum foxii** (Robertson, 1895) | | 1 |
| *Lasioglossum illinoense* (Robertson, 1892) | 4 | 2 |
| *Lasioglossum imitatum* (Smith, 1853) | 9 | 1 |
| *Lasioglossum laevissimum* (Smith, 1853) | 5 | 1 |
| *Lasioglossum leucozonium* (Schrank, 1791) | 98 | 17 |
| *Lasioglossum lineatulum* (Crawford, 1906) | 11 | 2 |
| *Lasioglossum nigroviride** (Graenicher, 1911) | | 3 |
| *Lasioglossum obscurum* (Robertson, 1892) | | 2 |
| *Lasioglossum oceanicum* (Cockerell, 1916) | 7 | |

| | | |
|---|---|---|
| *Lasioglossum paraforbesii* McGinley, 1986 | 3 | |
| *Lasioglossum pectorale* (Smith, 1853) | 197 | 5 |
| *Lasioglossum perpunctatum* (Ellis, 1913) | 23 | 1 |
| *Lasioglossum pilosum* (Smith, 1853) | 102 | 1 |
| *Lasioglossum subviridatum* (Cockerell, 1938) | | 1 |
| *Lasioglossum tegulare* (Robertson, 1890) | 4 | |
| *Lasioglossum timothyi* Gibbs, 2010 | | 8 |
| *Lasioglossum versans* (Lovell, 1905) | | 3 |
| *Lasioglossum versatum* (Robertson, 1902) | 1 | 13 |
| *Lasioglossum vierecki* (Crawford, 1904) | 67 | |
| *Sphecodes confertus* Say, 1837 | 1 | |
| *Sphecodes cressonii* (Robertson, 1903) | 3 | |
| *Sphecodes davisii* Robertson, 1897 | | 2 |
| *Sphecodes dichrous* Smith, 1853 | 10 | |
| *Sphecodes galerus* Lovell and Cockerell, 1907 | | 1 |
| *Sphecodes heraclei* Robertson, 1897 | 2 | 4 |
| *Sphecodes mandibularis* Cresson, 1872 | 10 | |

| | | | |
|---|---|---|---|
| | *Sphecodes ranunculi* Robertson, 1897 | 1 | |
| Megachilidae | *Coelioxys modesta* Smith, 1854 | 1 | |
| | *Coelioxys octodentata* Say, 1824 | 5 | |
| | *Coelioxys rufitarsis* Smith, 1854 | 7 | 1 |
| | *Heriades carinata* Cresson, 1864 | 45 | 2 |
| | *Heriades leavitti* Crawford, 1913 | 8 | 7 |
| | *Heriades variolosa* (Cresson, 1872) | | 6 |
| | *Hoplitis albifrons* (Kirby, 1837) | 1 | |
| | *Hoplitis pilosifrons* (Cresson, 1864) | 48 | |
| | *Hoplitis producta* (Cresson, 1864) | 1 | 1 |
| | *Hoplitis spoliata* (Provancher, 1864) | 3 | |
| | *Megachile addenda* Cresson, 1878 | 2 | |
| | *Megachile brevis* Say, 1837 | 9 | |
| | *Megachile campanulae* (Robertson, 1903) | 2 | 5 |
| | *Megachile gemula* Cresson, 1878 | | 1 |
| | *Megachile latimanus* Say, 1823 | 35 | 5 |
| | *Megachile mendica* Cresson, 1878 | 24 | 4 |
| | *Megachile montivaga* Cresson, 1878 | 4 | |

| | | |
|---|---|---|
| *Megachile pugnata* Say, 1837 | 9 | 5 |
| *Megachile relativa* Cresson, 1878 | 3 | 5 |
| *Osmia atriventris* Cresson, 1864 | 12 | 1 |
| *Osmia bucephala* Cresson, 1864 | | 4 |
| *Osmia cornifrons* (Radoszkowski, 1887) | | 4 |
| *Osmia distincta* Cresson, 1864 | 3 | |
| *Osmia georgica* Cresson, 1878 | 2 | 3 |
| *Osmia lignaria* Say, 1837 | 2 | |
| *Osmia pumila* Cresson, 1864 | 34 | 5 |
| *Osmia simillima* Smith, 1853 | 2 | |
| *Osmia texana* Cresson, 1872 | 5 | |

*Captured elsewhere in the E.S. George Reserve in 1972/1973 but not at “Evans’ Old Field”

**Table S4. Bee s**pecies change in relative abundance between the intensive historical sampling period (1972 and 1973) and the intensive contemporary sampling period (2017 and 2018) at the E.S. George Reserve. Only species that were captured in both sampling periods could be included in this analysis. Percent change calculations were then conducted on species with at least 30 records across the two sampling periods. Species were considered to be declining in abundance if they had >30% decrease in relative abundance, increasing if they had a >30% increase in relative abundance, and stable if they fell in between these categories.

| **Species** | **Historical Raw Abundance** | **Historical Relative Abundance** | **Contemporary Raw Abundance** | **Contemporary Relative Abundance** | **% Change** | **Change Classification** |
|---|---|---|---|---|---|---|
| *Agapostemon texanus* | 19 | 0.0106 | 1 | 0.0011 | | |
| *Andrena carlini* | 1 | 0.0006 | 1 | 0.0011 | | |
| *Andrena crataegi* | 61 | 0.0341 | 3 | 0.0032 | -90.6 | **Decreasing** |
| *Andrena forbesii* | 10 | 0.0056 | 9 | 0.0096 | | |

| | | | | | | |
|---|---|---|---|---|---|---|
| *Andrena hirticincta* | 19 | 0.0106 | 7 | 0.0075 | | |
| *Andrena nivalis* | 1 | 0.0006 | 1 | 0.0011 | | |
| *Andrena nubecula* | 12 | 0.0067 | 19 | 0.0202 | 201.7 | **Increasing** |
| *Andrena platyparia* | 1 | 0.0006 | 1 | 0.0011 | | |
| *Andrena rugosa* | 3 | 0.0017 | 3 | 0.0032 | | |
| *Andrena vicina* | 32 | 0.0179 | 6 | 0.0064 | -64.3 | **Decreasing** |
| *Anthophora terminalis* | 3 | 0.0017 | 1 | 0.0011 | | |
| *Augochlora pura* | 7 | 0.0039 | 111 | 0.1182 | 2921.1 | **Increasing** |

| | | | | | | |
|---|---|---|---|---|---|---|
| *Augochlorella aurata* | 126 | 0.0704 | 48 | 0.0511 | -27.4 | **Stable** |
| *Augochloropsis metallica* | 37 | 0.0207 | 13 | 0.0138 | -33.1 | **Decreasing** |
| *Bombus bimaculatus* | 95 | 0.0531 | 69 | 0.0735 | 38.4 | **Increasing** |
| *Bombus citrinus* | 3 | 0.0017 | 25 | 0.0266 | | |
| *Bombus griseocollis* | 19 | 0.0106 | 9 | 0.0096 | | |
| *Bombus impatiens* | 29 | 0.0162 | 365 | 0.3887 | 2297.9 | **Increasing** |
| *Bombus perplexus* | 8 | 0.0045 | 4 | 0.0043 | | |
| *Bombus vagans* | 33 | 0.0184 | 16 | 0.0170 | -7.6 | **Stable** |

| | | | | | | |
|---|---|---|---|---|---|---|
| *Ceratina dupla* | 166 | 0.0928 | 2 | 0.0021 | -97.7 | **Decreasing** |
| *Coelioxys rufitarsis* | 7 | 0.0039 | 1 | 0.0011 | | |
| *Colletes simulans* | 37 | 0.0207 | 2 | 0.0021 | -89.7 | **Decreasing** |
| *Dufourea monardae* | 31 | 0.0173 | 47 | 0.0501 | 188.9 | **Increasing** |
| *Epeolus scutellaris* | 15 | 0.0084 | 3 | 0.0032 | | |
| *Halictus confusus* | 89 | 0.0497 | 1 | 0.0011 | -97.9 | **Decreasing** |
| *Halictus ligatus* | 106 | 0.0593 | 45 | 0.0479 | -19.1 | **Stable** |
| *Heriades carinata* | 45 | 0.0252 | 2 | 0.0021 | -91.5 | **Decreasing** |

| | | | | | | |
|---|---|---|---|---|---|---|
| *Heriades leavitti* | 8 | 0.0045 | 7 | 0.0075 | | |
| *Hoplitis producta* | 1 | 0.0006 | 1 | 0.0011 | | |
| *Hylaeus affinis* | 55 | 0.0307 | 1 | 0.0011 | -96.5 | **Decreasing** |
| *Hylaeus mesillae* | 28 | 0.0157 | 1 | 0.0011 | | |
| *Hylaeus modestus* | 9 | 0.0050 | 2 | 0.0021 | | |
| *Lasioglossum anomalum* | 9 | 0.0050 | 6 | 0.0064 | | |
| *Lasioglossum coriaceum* | 42 | 0.0235 | 7 | 0.0075 | -68.2 | **Decreasing** |
| *Lasioglossum cressonii* | 13 | 0.0073 | 11 | 0.0117 | | |

| | | | | | | |
|---|---|---|---|---|---|---|
| *Lasioglossum illinoense* | 4 | 0.0022 | 2 | 0.0021 | | |
| *Lasioglossum imitatum* | 9 | 0.0050 | 1 | 0.0011 | | |
| *Lasioglossum laevissimum* | 5 | 0.0028 | 1 | 0.0011 | | |
| *Lasioglossum leucozonium* | 98 | 0.0548 | 17 | 0.0181 | -67.0 | **Decreasing** |
| *Lasioglossum lineatulum* | 11 | 0.0061 | 2 | 0.0021 | | |
| *Lasioglossum pectorale* | 197 | 0.1101 | 5 | 0.0053 | -95.2 | **Decreasing** |
| *Lasioglossum perpunctatum* | 23 | 0.0129 | 1 | 0.0011 | | |

| | | | | | | |
|---|---|---|---|---|---|---|
| *Lasioglossum pilosum* | 102 | 0.0570 | 1 | 0.0011 | -98.1 | **Decreasing** |
| *Lasioglossum versatum* | 1 | 0.0006 | 13 | 0.0138 | | |
| *Megachile campanulae* | 2 | 0.0011 | 5 | 0.0053 | | |
| *Megachile latimanus* | 35 | 0.0196 | 5 | 0.0053 | -72.8 | **Decreasing** |
| *Megachile mendica* | 24 | 0.0134 | 4 | 0.0043 | | |
| *Megachile pugnata* | 9 | 0.0050 | 5 | 0.0053 | | |
| *Megachile relativa* | 3 | 0.0017 | 5 | 0.0053 | | |

| | | | | | | |
|---|---|---|---|---|---|---|
| *Melissodes tinctus* | 4 | 0.0022 | 1 | 0.0011 | | |
| *Nomada cressonii* | 25 | 0.0140 | 4 | 0.0043 | | |
| *Nomada maculata* | 7 | 0.0039 | 3 | 0.0032 | | |
| *Osmia atriventris* | 12 | 0.0067 | 1 | 0.0011 | | |
| *Osmia georgica* | 2 | 0.0011 | 3 | 0.0032 | | |
| *Osmia pumila* | 34 | 0.0190 | 5 | 0.0053 | -72.0 | **Decreasing** |
| *Sphecodes heraclei* | 2 | 0.0011 | 4 | 0.0043 | | |

**Table S5.** Non-linear estimates of change over time (GAMs) for bee species collected in the ESGR with more than 20 specimens. Only those where year was significant are included.

| **Species** | **k'** | **Def. Expl** | **R2(adj)** | **edf** | **F** | **Ref. df** | **n** | **p** |
|---|---|---|---|---|---|---|---|---|
| *Andrena ceanothi* | 9 | 52.10% | 0.351 | 6.04 | 1.792 | 9 | 24 | 0.046 |
| *Augochlora pura* | 9 | 57.10% | 0.539 | 1.59 | 2.981 | 9 | 24 | < 0.001 |
| *Augochloropsis metallica* | 9 | 63.50% | 0.468 | 7.23 | 2.797 | 9 | 24 | 0.016 |
| *Bombus impatiens* | 9 | 79.50% | 0.771 | 2.44 | 8.412 | 9 | 24 | < 0.001 |
| *Ceratina calcarata* | 9 | 50.30% | 0.459 | 1.85 | 2.212 | 9 | 24 | < 0.001 |
| *Ceratina dupla* | 9 | 63.50% | 0.454 | 7.64 | 2.682 | 9 | 24 | 0.025 |
| *Colletes americanus* | 9 | 64% | 0.518 | 5.81 | 3.169 | 9 | 24 | 0.004 |
| *Halictus ligatus* | 9 | 29.60% | 0.264 | 0.985 | 0.858 | 9 | 24 | 0.006 |
| *Hylaeus affinis* | 10 | 41.30% | 0.316 | 3.25 | 2.93 | 3.863 | 24 | 0.04 |
| *Lasioglossum lineatulum* | 9 | 47.90% | 0.418 | 2.42 | 1.946 | 9 | 24 | 0.002 |
| *Lasioglossum pilosum* | 9 | 79.60% | 0.738 | 5.05 | 7.487 | 9 | 24 | < 0.001 |
| *Lasioglossum perpunctatum* | 9 | 19.60% | 0.161 | 0.967 | 0.485 | 9 | 24 | 0.033 |
| *Megachile pugnata* | 9 | 61% | 0.487 | 5.47 | 2.765 | 9 | 24 | 0.005 |

**Table S6.** Last year of collection for all species recorded at the E.S. George Reserve. No known collections at the ESGR occurred between 1999 and 2017.

| Species | Last year collected |
|---|---|
| *Agapostemon sericeus* | 1989 |
| *Agapostemon splendens* | 1987 |
| *Agapostemon texanus* | 2017 |
| *Agapostemon virescens* | 1989 |
| *Andrena aliciae* | 1977 |
| *Andrena alleghaniensis* | 1984 |
| *Andrena barbilabris* | 2018 |
| *Andrena bradleyi* | 1981 |
| *Andrena brevipalpis* | 2017 |
| *Andrena canadensis* | 1987 |
| *Andrena carlini* | 2018 |
| *Andrena carolina* | 1989 |
| *Andrena ceanothi* | 1987 |
| *Andrena commoda* | 1972 |
| *Andrena crataegi* | 2017 |
| *Andrena cressonii* | 1973 |
| *Andrena distans* | 1983 |
| *Andrena dunningi* | 2018 |
| *Andrena erigeniae* | 1972 |
| *Andrena forbesii* | 2018 |
| *Andrena fragilis* | 1973 |
| *Andrena hippotes* | 1971 |
| *Andrena hirticincta* | 2017 |
| *Andrena illinoensis* | 1937 |
| *Andrena imitatrix* | 1989 |
| *Andrena krigiana* | 2018 |
| *Andrena mandibularis* | 1980 |
| *Andrena mariae* | 1978 |
| *Andrena melanochroa* | 1980 |
| *Andrena miranda* | 1983 |
| *Andrena miserabilis* | 2018 |
| *Andrena nasonii* | 1984 |
| *Andrena nivalis* | 2017 |
| *Andrena nubecula* | 2017 |
| *Andrena perplexa* | 2018 |
| *Andrena placata* | 1985 |
| *Andrena platyparia* | 2017 |
| *Andrena robertsonii* | 1957 |
| *Andrena rudbeckiae* | 1982 |
| *Andrena rugosa* | 2017 |
| *Andrena salictaria* | 1978 |
| *Andrena sigmundi* | 1983 |
| *Andrena vicina* | 2018 |
| *Andrena virginiana* | 1987 |
| *Andrena wilkella* | 1989 |
| *Anthidiellum notatum* | 1975 |
| *Anthidium psoraleae* | 1983 |
| *Anthophora abrupta* | 1976 |
| *Anthophora terminalis* | 2018 |
| *Augochlora pura* | 2018 |
| *Augochlorella aurata* | 2018 |
| *Augochlorella persimilis* | 2018 |
| *Augochlorella pura* | 1999 |
| *Augochloropsis metallica* | 2018 |
| *Bombus affinis* | 1987 |
| *Bombus ashtoni* | 1989 |
| *Bombus auricomus* | 1975 |
| *Bombus bimaculatus* | 1987 |
| *Bombus borealis* | 1980 |
| *Bombus citrinus* | 2018 |
| *Bombus fernaldae* | 1972 |
| *Bombus fervidus* | 1987 |
| *Bombus frigidus* | 1960 |
| *Bombus griseocollis* | 2018 |
| *Bombus impatiens* | 2018 |
| *Bombus pensylvanicus* | 1982 |
| *Bombus perplexus* | 2017 |
| *Bombus terricola* | 1978 |
| *Bombus vagans* | 2018 |
| *Calliopsis andreniformis* | 1982 |
| *Calliopsis nebraskensis* | 1983 |
| *Ceratina calcarata* | 2018 |
| *Ceratina dupla* | 2018 |
| *Ceratina mikmaqi* | 2018 |
| *Ceratina strenua* | 2018 |
| *Chelostoma philadelphi* | 1989 |
| *Coelioxys alternatus* | 1987 |
| *Coelioxys immaculatus* | 1960 |
| *Coelioxys moestus* | 1972 |
| *Coelioxys octodentatus* | 1983 |

| | |
|---|---|
| *Coelioxys rufitarsis* | 2017 |
| *Colletes americanus* | 1980 |
| *Colletes compactus* | 1975 |
| *Colletes inaequalis* | 1989 |
| *Colletes kincaidii* | 1977 |
| *Colletes latitarsis* | 1977 |
| *Colletes nudus* | 1973 |
| *Colletes simulans* | 2017 |
| *Colletes solidaginis* | 1987 |
| *Colletes validus* | 1989 |
| *Dufourea monardae* | 2018 |
| *Epeolus ainsliei* | 1960 |
| *Epeolus bifasciatus* | 1982 |
| *Epeolus canadensis* | 1989 |
| *Epeolus interruptus* | 1982 |
| *Epeolus lectoides* | 1987 |
| *Epeolus minimus* | 1975 |
| *Epeolus pusillus* | 1985 |
| *Epeolus scutellaris* | 2017 |
| *Halictus confusus* | 2017 |
| *Halictus ligatus* | 2018 |
| *Halictus parallelus* | 1989 |
| *Halictus rubicundus* | 1989 |
| *Heriades carinata* | 2017 |
| *Heriades leavitti* | 2018 |
| *Heriades variolosa* | 2018 |
| *Holcopasites calliopsidis* | 1973 |
| *Hoplitis pilosifrons* | 1989 |
| *Hoplitis producta* | 2018 |
| *Hoplitis spoliata* | 1980 |
| *Hylaeus affinis* | 2017 |
| *Hylaeus annulatus* | 1974 |
| *Hylaeus illinoisensis* | 2017 |
| *Hylaeus mesillae* | 2017 |
| *Hylaeus modestus* | 2018 |
| *Hylaeus sp. A* | 2017 |
| *Lasioglossum imitatum* | 2017 |
| *Lasioglossum abanci* | 2017 |
| *Lasioglossum admirandum* | 1957 |
| *Lasioglossum anomalum* | 2018 |
| *Lasioglossum bruneri* | 1985 |
| *Lasioglossum cattellae* | 2018 |
| *Lasioglossum cinctipes* | 1978 |
| *Lasioglossum coeruleum* | 2018 |
| *Lasioglossum coriaceum* | 2018 |
| *Lasioglossum cressonii* | 2018 |
| *Lasioglossum ellisiae* | 1960 |
| *Lasioglossum forbesii* | 1976 |
| *Lasioglossum foveolatum* | 1987 |
| *Lasioglossum foxii* | 2018 |
| *Lasioglossum illinoense* | 2017 |
| *Lasioglossum imitatum* | 1987 |
| *Lasioglossum laevissimum* | 2017 |
| *Lasioglossum leucocomum* | 2018 |
| *Lasioglossum leucozonium* | 2018 |
| *Lasioglossum lineatulum* | 2017 |
| *Lasioglossum lustrans* | 1983 |
| *Lasioglossum nelumbonis* | 1978 |
| *Lasioglossum nigroviride* | 2017 |
| *Lasioglossum nymphaearum* | 1961 |
| *Lasioglossum obscurum* | 2018 |
| *Lasioglossum oceanicum* | 1977 |
| *Lasioglossum paraforbesii* | 1973 |
| *Lasioglossum pectorale* | 2018 |
| *Lasioglossum perpunctatum* | 2017 |
| *Lasioglossum pictum* | 1961 |
| *Lasioglossum pilosum* | 2017 |
| *Lasioglossum planatum* | 1983 |
| *Lasioglossum smilacinae* | 1990 |
| *Lasioglossum subviridatum* | 2018 |
| *Lasioglossum tegulare* | 1979 |
| *Lasioglossum texanum* | 1949 |
| *Lasioglossum timothyi* | 2018 |
| *Lasioglossum versans* | 2018 |
| *Lasioglossum versatum* | 2017 |
| *Lasioglossum vierecki* | 1999 |
| *Megachile addenda* | 1985 |
| *Megachile brevis* | 1990 |
| *Megachile campanulae* | 2018 |
| *Megachile centuncularis* | 1974 |
| *Megachile frugalis* | 1979 |
| *Megachile gemula* | 2018 |
| *Megachile latimanus* | 2018 |

| | |
|---|---|
| *Megachile mendica* | 2018 |
| *Megachile pugnata* | 2018 |
| *Megachile relativa* | 2018 |
| *Melissodes apicatus* | 1983 |
| *Melissodes bimaculatus* | 2018 |
| *Melissodes communis* | 1974 |
| *Melissodes dentiventris* | 1963 |
| *Melissodes desponsus* | 2018 |
| *Melissodes druriellus* | 1978 |
| *Melissodes iliatus* | 2018 |
| *Melissodes illatus* | 2018 |
| *Melissodes niveus* | 1987 |
| *Melissodes subillatus* | 1989 |
| *Melissodes tinctus* | 2017 |
| *Nomada armatella* | 1973 |
| *Nomada articulata* | 1986 |
| *Nomada cressonii* | 2018 |
| *Nomada cuneata* | 2018 |
| *Nomada dreisbachi* | 1978 |
| *Nomada fervida* | 1960 |
| *Nomada lepida* | 1978 |
| *Nomada maculata* | 2018 |
| *Nomada ovata* | 1984 |
| *Nomada perplexa* | 1974 |
| *Nomada pygmaea* | 1983 |
| *Nomada rubicunda* | 1984 |
| *Nomada sayi* | 1972 |
| *Nomada subrutila* | 1989 |
| *Nomada sulphurata* | 1973 |
| *Nomada vicina* | 1985 |

| | |
|---|---|
| *Osmia albiventris* | 1989 |
| *Osmia atriventris* | 2018 |
| *Osmia bucephala* | 2018 |
| *Osmia caerulescens* | 1943 |
| *Osmia cornifrons* | 2018 |
| *Osmia distincta* | 1978 |
| *Osmia georgica* | 2017 |
| *Osmia pumila* | 2018 |
| *Osmia simillima* | 1972 |
| *Osmia texana* | 1977 |
| *Peponapis pruinosa* | 2017 |
| *Perdita bequaerti* | 1972 |
| *Perdita octomaculata* | 1972 |
| *Pseudopanurgus aestivalis* | 1978 |
| *Pseudopanurgus andrenoides* | 2017 |
| *Sphecodes aff. atlantis* | 1989 |
| *Sphecodes confertus* | 1984 |
| *Sphecodes cressonii* | 1982 |
| *Sphecodes davisii* | 2017 |
| *Sphecodes dichrous* | 1986 |
| *Sphecodes galerus* | 2017 |
| *Sphecodes heraclei* | 2017 |
| *Sphecodes illinoensis* | 1977 |
| *Sphecodes mandibularis* | 1990 |
| *Sphecodes ranunculi* | 1972 |
| *Triepeolus donatus* | 1972 |
| *Triepeolus simplex* | 1982 |
| *Xylocopa virginica* | 2017 |

**Table S7:** Summary stats for GLMM on phenological range across bee categories from Fig 2e. Phenological ranges measure the flight period phenology of bee species measured in Julian days. Longer phenological ranges indicate longer flight periods and vice versa. Neural network analysis revealed a consistently important, but qualitatively different effect of the phenological range variable on the probability of extinction across different categories of bees (see Results). We used a GLMM (glmer R package) to verify and visualize the result in Figure 2e.

| GLMM: Extinction Probability ~ ( 1 + phenological range \| bee type)<br>Family: binomial (link = logit) | | | |
|---|---|---|---|
| AIC | Log Likelihood | Deviance | Df.residual |
| 177.4 | -84.7 | 169.4 | 132 |
| Random Effects (n=136 species across 5 bee types) | | | |
| Group | Name | Variance | Std. Dev. |
| bee type | Intercept | 1.611 | 1.270 |
| | phenological range | 0.001 | 0.032 |
| Fixed Effects | | | |
| | Estimate | Std. Error. | p value |
| Intercept | 0.866 | 0.293 | 0.003 |

**Open data access:**

Data used for analyses in this manuscript, including Evans' original dataset from 1972/1973 with updated species nomenclature, will be permanently archived at the USDA Ag Data Commons after the acceptance of this manuscript and will be citable and accessible here: https://data.nal.usda.gov/dataset/century-sampling-ecological-preserve-reveals-declining-diversity-wild-bees.

University of Michigan Museum of Zoology Insect Collection data are available for download at: https://www.gbif.org/dataset/13e7869e-0c76-473a-a227-53d6e3d6fbf2. Data are also available at https://scan-bugs.org/portal/collections/harvestparams.php. Inclusion of records collected or databased as part of this project will be added to these databases following the acceptance of this manuscript for publication. Use the Locality search term: "E.S. George Reserve" to find all records for the reserve.

Intertegular Distance data and bee traits data were sourced from published articles or datasets. Data and sources of data, including DOI and/or website links for the data will be available on the USDA Ag Data Commons dataset (see above). The trait file is also available on DataDryad for review: https://datadryad.org/stash/share/TVZGtBbRd3x1lt9x__VM5ZzRfBIEO1JgkKBWyzRsb38.

We downloaded data available on GBIF. To determine the latitudinal and longitudinal ranges of species we did not include any limits other than a cutoff of 20,000 records, to reduce data processing time. For the phenological search, we also included a cutoff of 20,000 records and included longitudinal and latitudinal bounds to ensure that the phenology data was relevant to Michigan bees. We therefore limited the search to specimens that appear within the bounds of -109< longitude <-70 and 37< latitude <48.

### Supplemental Methods

#### Sampling and Records Gathering

***Contemporary sampling* -** In 2017, the first sampling day was June 1 and the final sampling day was September 25. In 2018, the first sampling day was May 8 and the final day was October 3. We expanded our sampling in 2018 to include a wider diversity of bees with narrower phenological periods. During each visit we sampled bees using three methods. First, we walked to the center of the open field and randomly selected a direction to start the first 25 meter transect. Three other 25 m transects were then established based off the first one, each at a 90 degree angle from the neighboring transect for a total of 100m sampled, with each transect segment moving away from a central location. Each transect was walked for 10 minutes each, a total of 40 minutes of sampling. We used aerial insect nets to collect bees found within 1.5m of the transect, and time was stopped for specimen processing. Second, we spent 20 minutes collecting bees from any plants in the general vicinity of the open field. Third, to most closely match the methods used by Evans, we spent 30 minutes sampling bees at each of the primary blooming plant species located in the field. This sampling method was always done last, and included any plants that we collected more than one bee from that day. All bees were identified to species (or lowest possible taxonomic level) using relevant keys (Mitchell 1960, 1962, Ribble 1967, LaBerge and Bouseman 1970, LaBerge 1971, 1973, 1977, 1980, 1985, 1989, Bouseman and LaBerge 1978, McGinley 1986, Coelho 2004, Rehan and Sheffield 2011, Gibbs 2011, Gibbs et al. 2013). All specimens collected in 2017 and 2018 are currently held in the Isaacs Lab at Michigan State University (as of 2022), and will be deposited at the A.J. Cook Arthropod Collection at Michigan State University for long term inclusion in that collection.

***Databasing museum records* -** The University of Michigan Museum of Zoology Insect Collection (UMMZI), Ann Arbor, MI, holds over 4,000 bee specimens from the historical collections at the ESGR. As part of this study, we worked with the UMMZI to catalog and transcribe the label data from these specimens. Each was assigned a unique identifier and was imaged alongside its respective data labels using the museum's imaging stations. Specimen/label images allowed for transcription options off site as well as facilitated ease of checking any potential transcription typos or specimens likely misidentified. Historical label data were transcribed from images and shared with the UMMZI for inclusion in their Specify database. The UMMZI database publicly shares specimen records via IPT to GBIF, SCAN, and others with records in this study to be made accessible upon study completion. Records will be accessible at: https://www.gbif.org/dataset/13e7869e-0c76-473a-a227-53d6e3d6fbf2 and https://scan-bugs.org/portal/collections/harvestparams.php.

Historical data were checked for entry errors and outdated taxonomies. Specimens with questionable species determinations were re-examined and re-identified using relevant keys (see above) where possible. When additional taxonomic expertise was required for identification and images did not suffice, specimens were loaned to the Gibbs Lab (University of Manitoba; current author) for identification. Bees that could not be confidently identified to the species level were excluded from the dataset, and entries that were missing the date of collection were also removed. Excluded entries accounted for less than 1% of the specimens.

In addition to analyses of the 4,000 plus records from the ESGR since 1921, we also conducted more focused analyses comparing Evans' dataset from his 1972 and 1973 collection effort to our contemporary collections in 2017 and 2018. Evans' original dataset from 1972/1973 was available through University of Michigan records, and we used these for our focused analyses of these years. The dataset is unique compared to the records from the museum, because Evans did not always collect observed bees if he was confident in their identification (especially *Bombus* spp. and oligolectic species, e.g., *Andrena rudbeckiae* and *Dufourea monardae*; Evans 1986). Therefore, his original dataset provides a more complete representation of the community he encountered.

### Expanded Methods for Data Analyses

*Community change across a century of data.* We employed an approach similar to that used by Bartomeus and colleagues (2013). To account for changes in practices across a century of records (1921-2018), such as including only a synoptic collection in the museum, we removed any duplicate specimens from this data. These were records of the same species collected on the same date (e.g., if there were five *Bombus impatiens* records for June 3, 1979, we only kept one of these records in the dataset for analyses) (see methods in Bartomeus et al. 2013). Then, to account for uneven sampling, we binned specimens into seven groupings of consecutive years to achieve more even sample numbers in each bin prior to comparison of species richness (also see Bartomeus et al. 2013) (Fig. S1). We then rarefied all bins to the bin with the lowest sample size (n = 320) using the rarefy function (package: vegan) to calculate the mean species richness and standard error of the mean. We used an ANOVA with a Tukey's means difference test to determine significant differences among the bins.

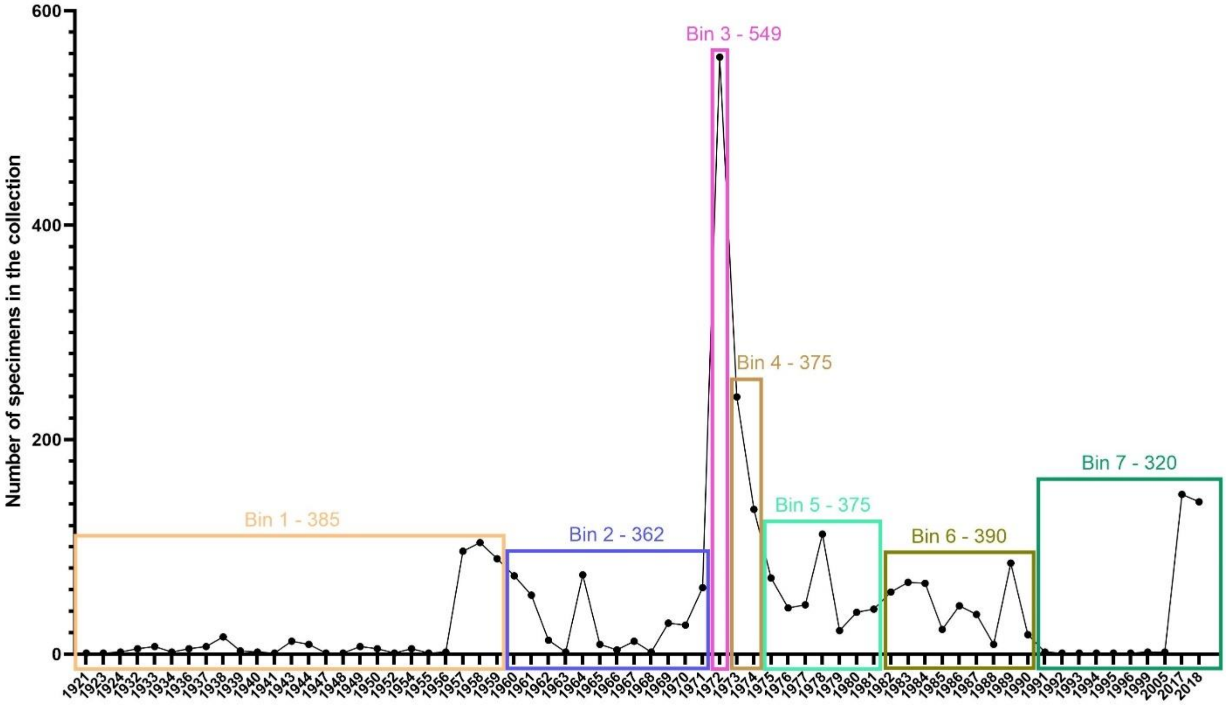


**Figure S1.** Number of bee specimen records from the E.S. George Reserve in each year after duplicate specimens within the same collection event were removed. The seven bin time periods are shown with the corresponding number of specimens in each bin used for comparison of bee species richness and community composition over time.

Using these processed data to account for changes in sampling practices and uneven sampling, we also estimated changes in relative abundance for a subset of species. To limit effects of low sample years, we only considered years with at least 30 observations (24 years). We also only included species with at least 20 records. We then calculated relative abundance in each sampling year (including years when sampling occurred but the species was not captured) by dividing the number of records for that species by the entire count of specimens documented that year (after removal of repeats). This helps to account for the temporal changes in sampling effort. We then used generalized additive models (GAMs) (function: gam, package: mgcv; Wood 2004, 2011, 2022) to explore trends in abundance over time with year as a smoothing factor (e.g., gam(([Abundance of Sphecodes mandibularis]/[total number of specimens that year])

~s(Year, k = -1, bs = “cs”), data = Sphecodes_mandibularis). Residuals were checked using the function gam.check (package: mgcv). Predictions were displayed using the function plot (package: mgcv) with standard error of the mean.

In addition to using the data processing methods similar to Bartomeus and colleagues (2013), we also conducted analyses which did not remove repeat samples or group several years together. Instead, we used methods primarily aimed to reduce the impact of undersampled years. We accomplished this via rarefied species richness in each year to compare equal sample sets for richness. Rarefaction was done using the R package vegan. To limit effects of low sample years, we only included years with 20, 30, 40, 50, 60, or 70 observations. Due to the apparent quadratic patterns, richness was regressed across sample years again using GAMs and minimized generalized cross-validation (GCV) for fitting.

*Measuring change in community composition between the contemporary and historical sampling periods.* We compared the contemporary sampling (years 2017 and 2018) to Evans’ original dataset from 1972 and 1973. We chose these years to compare due to similarities in sampling methods and intensity. Though only data from months with sampling efforts in both the contemporary and historical data sets were included (May, June, July, August, September). We rarefied the species richness for the historical data and the contemporary data (function: rarefy, package: vegan) (Oksanen et al. 2019) to the lowest sample size (contemporary data: n = 1271). We compared mean (+/- SE) rarefied species richness between the historical and the contemporary sampling years using an unpaired two-tailed t-test. We also calculated Shannon diversity (Hill number) for each sampling effort (function: hill_div; package: hilldiv) (Alberdi 2019), which accounts for both species richness and evenness in the population. A higher diversity estimate would indicate high species richness and evenness, whereas lower diversity can be due to low richness or evenness or both. We rounded up to whole numbers to represent an estimate of the number of species (as Hill number's represent the effective number of species). Shannon diversity was statistically compared between datasets using Hutcheson t-test for two communities (function: Hutcheson_t_test, package: ecolTest) (Salinas 2021). Additionally, we calculated Pielou's evenness index ([Shannon diversity]/log(species richness) for both datasets.

Non-metric multidimensional scaling (NMDS) ordinations were used to visualize differences in species composition between the historical and contemporary datasets (vegan package). We then used permutational multivariate analysis of variance (PERMANOVA) (function: adonis, package: vegan) (Anderson 2001) to determine whether the bee communities were significantly dissimilar in ordination between the two sampling periods. Since PERMANOVA is sensitive to differences in dispersion among groups, regardless of ordination, we first tested for homogeneity of dispersion using the betadisper function (package: vegan). This is a multivariate analog of Levene's test for homogeneity of variances that implements PERMDISP2 (Anderson 2004). Homogeneity of dispersion paired with significant differences between groups according to the PERMANOVA would therefore indicate that the community composition differed between historical and contemporary samples. A dummy species was added to the species matrix prior to analyses and analyses were based on Bray-Curtis dissimilarity, with 999 permutations. We also used the envfit function (package: vegan) to determine which members of the bee community were significantly ($p < 0.01$) contributing to ordination in the NMDS. We fit vectors on the NMDS plot according to species contributions, with the length of the vector corresponding to the strength of the contribution.

To compare dominant species within the two communities, we calculated the percent composition of each species to the overall community in the historical sampling years and the contemporary years. These values were compared for species that made up over 2%.

We also calculated the percent change in relative abundance of species that were collected in both the historical and contemporary samples. To calculate relative abundance, we summed the number of records for each species collected in each sampling period (records were summed for 1972/1973 and for 2017/2018). We then calculated the relative abundance by dividing the summed species record by the total number of specimens in that collection period. This was to account for more sampling effort (more days sampling and more specimens collected) in the historical period. We then calculated the percent change in relative abundance for species with at least 30 records across the two sampling periods. Species were designated as declining if they had an abundance decline of more than 30% and increasing if they had an increase of more than 30%. Species in between were considered stable. This cutoff was chosen to match the IUCN designation for vulnerable species (IUCN Standards and Petitions Subcommittee 2014).

*Measuring change in community traits between contemporary and historical sampling periods.* Species natural history characteristics and traits were collected and collated for the species in our study (Table S1). We include taxonomic designation from family to species. Nest location, sociality, and dietary breadth were listed categorically and determined based on literature review and expert knowledge for as many species as possible. We established latitudinal and longitudinal ranges per species with specimen location data available on the Global Biodiversity Information Facility (GBIF), using a 20,000 sample cutoff for the most numerous entries (e.g.,

*Bombus impatiens*). Phenological ranges of bee flight periods were also sourced via GBIF sample dates (measured in Julian days) but with latitudinal and longitudinal limits to ensure proximity to our sample site. Phenological data per species were downloaded from GBIF by searching for specimen data within the bounds of -109< longitude <-70 and 37< latitude <48 to limit phenological data to regions heuristically similar to the ESGR. We sourced intertegular distance values (ITD; mm) from published data (Gibbs et al. n.d., Castillo and Fairbairn 2012, Bartomeus et al. 2013, Cariveau et al. 2016, Lerman and Milam 2016, Hung 2017, Normandin et al. 2017, Nicholson 2019, Kendall et al. 2019, Lim et al. 2022) and specimen measurements conducted at the J. B. Wallis / R. E. Roughley Museum of Entomology at the University of Manitoba by J. Gibbs for species where published ITD data were not available (Table S1). Three species for which ITD could not be sourced in the literature (*Melissodes niveus, M. tinctus,* and *Nomada subrutila*) or conducted at the J. B. Wallis / R. E. Roughley Museum of Entomology were filled in using means of the ITD for other species in the same genus. Removal of these three species from analysis did not qualitatively change results. ITD data were then used to calculate flight distance (max homing dist; km; Greenleaf et al. 2007) and proboscis length (mm; Kendall et al. 2019).

Trait data were used in two types of analyses: 1) At the species level we used traits as potential predictors of species persistence or local extirpation in the ESGR between the historical and contemporary periods, and 2) At the community level, we compared trait composition between the two periods' sample communities.

In using species traits as predictors of persistence between periods, we first assigned all species present in the historical dataset with an additional binary variable "extirpation" indicating if the population persisted into the contemporary sampling period or is locally extirpation (not collected in 2017-2018). This extirpation variable functioned as our predicted variable. Given the mixed trait data set (both categorical & continuous trait variables), we initially pursued the use of mixed linear models to determine if certain traits are associated with local extirpation. However, non-linear relationships (see Results/Supplementry Results) and highly linked categorical variables between predictors (e.g., eusociality linked with polylecty) limited the suitability and interpretability of linear models for these data. We also attempted to garner insight through ordination methods comparing the traits of extirpation versus persistent species. However, little variance could be explained via permANOVA and NMDS (see Results). Additionally, PCA analysis was limited by the high dimensionality required to explain the variance in our data and the number of predictors which highly contributed to multiple principal components across numerous subsets of our predictors (see Fig S2).

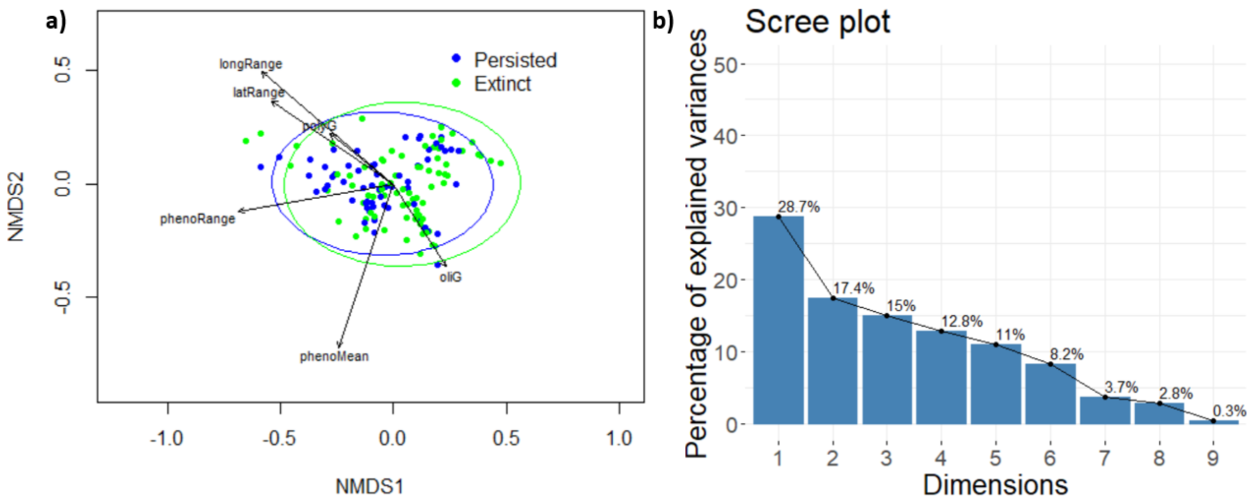


**Figure S2.** Ordination analysis between sample periods at species level for bees collected at the E.S. George Reserve. a) NMDS, accompanying permANOVA indicates significant difference (p=0.012), but can only explain 3% of variance (F_1,134=4.00). b) Scree plot detailing results of example PCA analysis. The PCA can explain full variance, but requires 6 dimensions to explain >90%. Additionally, 3+ variables contribute to multiple dimensions (including the most explanatory). In both a) and b), these qualities substantially limit any possible inference garnered from these more traditional techniques. Note, both of the presented analyses here were completed with quantitative traits: phenological mean, phenological range, latitudinal range, longitudinal range, ITD and categorical traits: oligolectic ground nesting (oliG) polylectic ground nesting (polyG), polylectic cavity nesting (polyC), and cleptoparasitic (clepto). Possible inference and general results were qualitatively unchanged across different tested subsets of trait variables.

We pursued neural networks due to their flexible non-parametric nature, ability to handle both categorical and continuous data, and increased capability to capture variance and feature effects despite lingering collinearity and limited data respectively. Species persistence from historical ('72/'73) to contemporary ('17/'18) sampling periods served as our dependent predicted variable in our neural network set up (see Fig S3). Prior to fitting neural networks, predictor variables (aka "features") were scaled using the scale package in R (Wickham and Seidel 2022). Categorical variables were expanded to new single factor columns using one-hot encoding (aka one-of-K scheme; mltools package; Gorman 2018). Further data preprocessing was done by removing redundant data via the cor function in R. Specifically, the regression equations used to predict proboscis length (mm) and flight distance (max homing dist; km) made each variable redundant to ITD so we only used ITD. Furthermore, both species' latitudinal and longitudinal ranges were highly correlated with both latitudinal and longitudinal maximums and minimums. However, latitudinal and longitudinal maximums and minimums showed no correlation between themselves or with each other. We therefore removed latitudinal and longitudinal ranges as redundant. Next, 10th and 90th phenology quantiles were strongly correlated with phenological means and slightly correlated with each other. We therefore removed the quantile variables from model based analysis. Finally, our categorical trait variables (nesting location, dietary range, socialiy) presented collinearity issues because 1) all shared the "clepoparasitic" class across variables and 2) some classes nearly predicted each other (e.g., eusociality always predicted polylectic diets). We therefore reformatted unique combinations as separate binary variables, e.g., oligolectic ground nesting species were marked as such in the new ‘oligolectic-ground’ variable.

We used shallow (single layer) neural networks (nnet R package; Venables and Ripley 2002), as these function best when data are limited for each class targeted for prediction (Basha et al. 2020). This also allowed us to use the caret package (Kuhn 2015) to train our classification task neural networks with cross validation. Models were trained on the whole data set (data from the historical and contemporary sampling periods). To limit over-fitting in trained models, the number of neurons was limited following Heaton (2008), providing comparable model error, performance (i.e., >95% accuracy), and predictor analysis (see results) with 3-6 neurons in our hidden layer with a decay rate during training ranging between 5e-2 - 5e-4. See Figure S3 for a visualized example network used in our analysis.

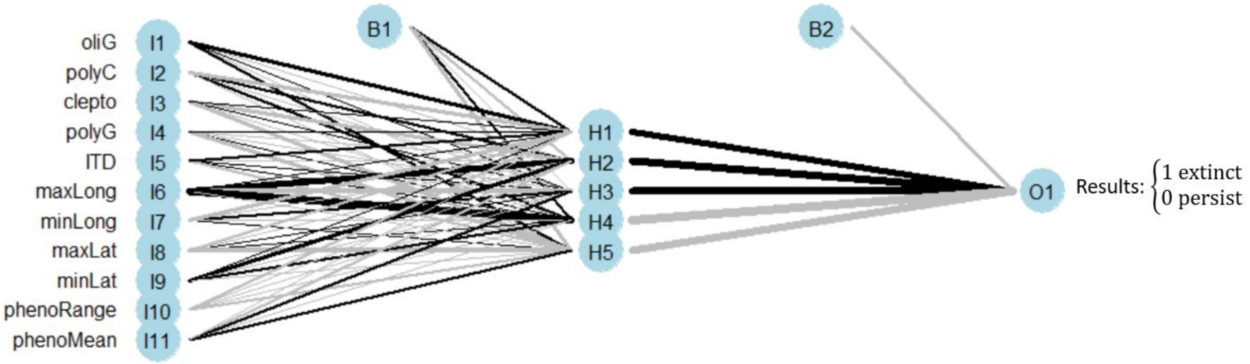


**Figure S3.** Example neural network trained on bee species persistence data from the ‘72-’73 sample period to ‘17-’18 sample period. 11 inputs (our prepared trait variables, see Methods) into a single hidden layer of 5 neurons with one output layer (binary extinct or persist variable). B1 and B2 neurons represent the network’s bias nodes. Black and gray lines represent positive and negative weights respectively with line thickness indicating the value of the weight. Decay rate set to 0.01, 97.5% accuracy over species persistence data.

We estimated the non-directional importance of each trait (predictor) in our trained model/network for its relative influence on the model predictions using the Garson method

(NeuralNetTools package; Beck 2018). Garson values range from 0 (indicating no relative importance/contribution in producing predictions) to 1 (indicating the highest relative importance/contribution in influencing predictions). We then used the Olden method (NeuralNetTools package) to estimate the effect of each trait (predictor) on influencing specific model predictions. The Olden method produces continuous values with positive values indicating a positive relationship with the predicted variable (i.e., influencing an extinct neural net prediction) and negative values indicating a negative relationship with the predicted variable (i.e., indicating a persistent neural net prediction). Olden importance values were rescaled between -1 and 1 per individual neuralnet/model run. We note that Olden importance values are not regression slopes. They do not indicate the linear relationship between the predictor variable and the probability of extirpation. Instead, they represent relative degrees of the most consistently positive (as values go to 1) or negative (as values go to -1) weights associated with each predictor and the predicted variable. For example, a value of -1 indicates a predictor had the most negative association with predicting extirpation out of the tested predictors. A value closer to 0 can indicate no strong relation or nonlinear relationships. For further descriptions see Olden et al. 2004. We verified feature effects in our neural networks implied via Olden values using partial dependence plots (PD plots; iml package) (Molnar 2022). Finally, sufficiently strong effects were tested with linear frequentist statistics (see Supplementary Results).

We also directly compared community traits in the historical and contemporary communities. We first compared traits between periods using simple comparisons of the composition of categorical variables (e.g., nesting location) and ranges of quantitative traits (e.g., intertegular distance) per sample population. These were used to corroborate/vet potential signals found in our neural network analysis on the species and community levels. We also separated community composition via averages of each quantitative trait metric and percent make up of each categorical trait (e.g., nesting location or lecty) by collection period and month. We again visualized the trait composition differences between the historical and contemporary communities via NMDS (see methods described above) and used PERMANOVA to test for statistical differences after confirming no differences in dispersion.

### Supplemental Results

#### *Changes in community composition across a century of sampling*

We used GAMs to test for significant changes in species richness over time. In this dataset, we included all museum records without removing duplicate records of species collected on the same days. Instead, to minimize the impact of low sampling years, we only included years with at least 20 specimens. We also conducted the analyses with yearly sample minimums of 30, 40, 60, and 70 to see if this changed the overall data trends. Again, we see a unimodal pattern of species richness (Fig. S4). First, we see initial increases in richness, peaking in the early 1970s. Second, we see a transition to species loss after reaching a maximum in the 70s. Figure 7 used a minimum of 50 observations/year for inclusion in the analysis to limit effects of low sample years. Repeating this analysis with five other sample minimums (20, 30, 40, 60, & 70) presented no qualitative changes.

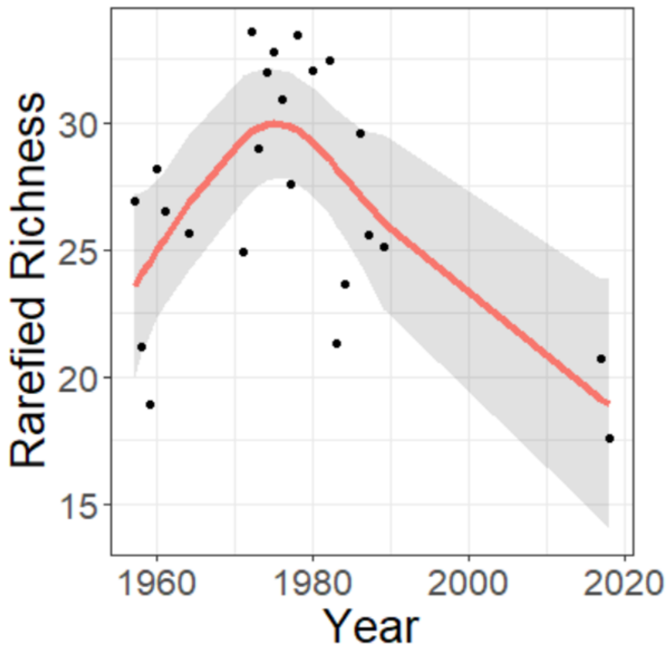


**Figure S4.** GAM regression of rarefied richness with a 50-sample minimum per year. F=5.755, edf=2.961, Ref.df=3.495, p=0.00402, Deviance explained = 54.1%, GCV = 14.868.

We also used GAMs to test for changes in relative abundance of common species. Year was significant for 13 species (Fig S5; Table S5), with varying trends over time. Some species had peaks in abundance around the 1970s and 1980s: *Andrena ceanothi, Ceratina dupla*, and *Hylaeus affinis* (Fig. S5). Others peaked in the 1960s and had an overall downward trend in abundance: *Colletes americanus*, *Lasioglossum lineatulum*, *L. perpunctatum*, and *L. pilosum* (Fig. S5). Others had generally upwards trends in abundance, peaking in the 2010s: *Augochlora pura*, *Bombus impatiens*, *Ceratina calcarata*, *Halictus ligatus* (Fig. S5). Others had more variable trends, such as *Augochloropsis metallica* which peaked in the 1980s and 2010s, and *Megachile pugnata*, which peaked in the 1990s and early 2000s (Fig. S5).

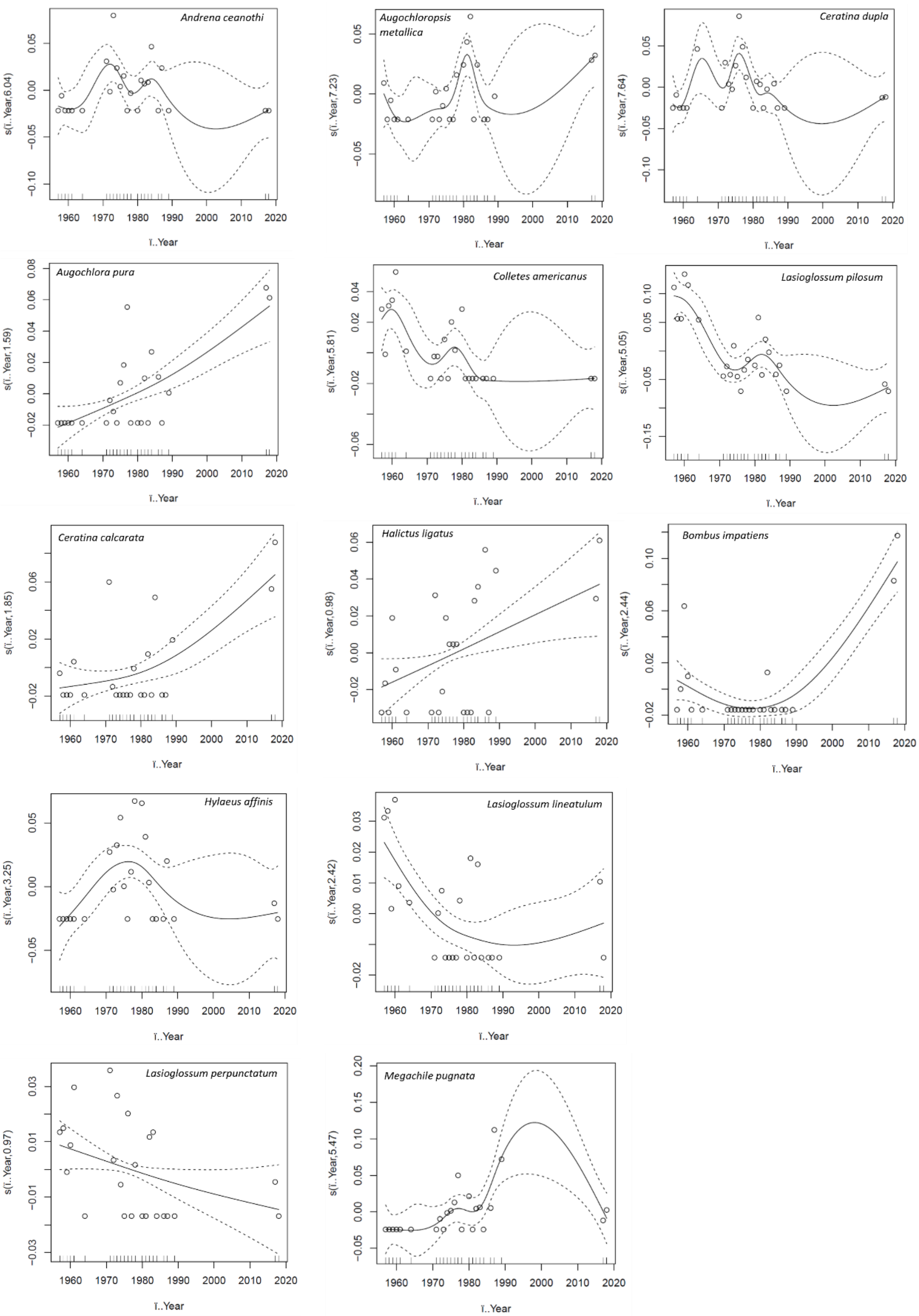


**Figure S5.** GAM regressions for bee species collected in the E.S. George Reserve with more than 20 records. Only those where year was significant are included. Summary stats available for all Fig S5 GAMs in Table S5.

***Community composition change between historical and contemporary sampling efforts***

The historical and contemporary bee communities did not differ significantly in level of dispersion (PERMDISP; $F_{1,9} = 1.49$, $p = 0.26$), and were significantly different from each other in community composition (PERMANOVA; $R^2 = 0.26$, $F_{1,9} = 2.87$, $p = 0.01$) (Fig. 1).

***Species traits as indicators of local extirpation***

Using traits to identify factors involved in species persistence between the historical and contemporary sampling periods had limited effect using traditional methods. A PERMANOVA analysis (Fig. S2a) indicated a significant trait-based difference between persistent and extirpated species, but the test only explained 3% of the variance. PCA analysis accounted for more variance, but the leading two principal components only explained 46% of the variance. A full six components are required to explain >90% of the variance, limiting any utility in reducing dimensionality with the ordination process (Fig. S2b).

On the other hand, a single trained neural network predicted species persistence with accuracy ranging from 82% on average when using three neurons to an average of 98% when using five neurons in our hidden layer (see example in Fig. S3). To estimate the influence of individual traits over network predictions, we trained 1000 iterations of our single layer networks with random initial weights to analyze the consistency of each trait's Olden importance and effect on predicted outcome. This resulted in some clear trends in our categorical variables (Fig. S6). Before highlighting our neural network results, we again note that Olden importance values are not necessarily akin to regression slopes or coefficient relationships between predictor and predicted variables. Instead, they represent the relative consistency of effect on predicting extirpation (1) or persistence (-1).

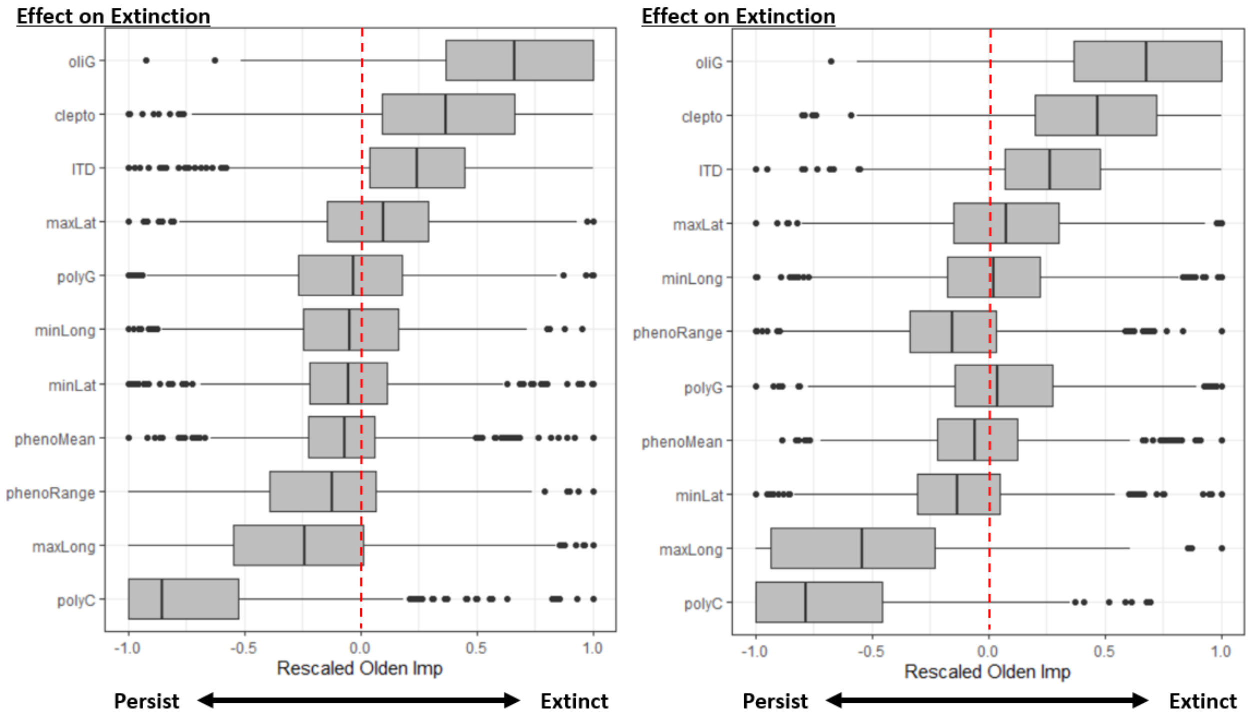


**Figure S6.** Summary of bee trait (predictor) importance regarding neural network predictions after 1000 training iterations on the full data set. a) Results show importance from models with a single hidden layer, 6 hidden neurons, and a decay rate of 5e-4. Importance was estimated using the Olden algorithm and box and whisker plots represent the range of 1000 iterations. Negative values indicate a negative relationship with the predicted variable (extinct, i.e., more likely to persist) while positive values indicate a positive relationship with the predicted variable (i.e., more likely to go extinct). The red line at 0 indicates no effect. Some predictors show clear trends in importance, but inference is limited in these models because all predictors exhibit a wide range of importance. Patterns are robust to network design as shown in example b) with 3 hidden neurons and a decay rate of 5e-2.

The most influential three variables were categorical; oligolectic ground nesting bees and cleptoparasitic bees were associated with higher probabilities of extirpation, while polylectic cavity nesting bees were most strongly linked to persistence (Fig. S6). These trends were corroborated by reviewing the percent persistence in each category in the raw data (Fig. 2a-d; Fig S7). However, further inference over our quantitative parameters was hindered by high levels of variability in quantitative trait effects on network predictions (see wide error bars in Fig. S6). This could indicate potential nonlinearities in these quantitative traits or distinct effects across different subsets of the data. Given the sharp distinctions in persistence across categorical traits

(Fig. 2), separate models were fit for each categorical subset in our data (see Supplementary Methods): 1) oligolectic ground nesting, 2) polylectic ground nesting, 3) polylectic cavity nesting, 4) cleptoparasitic. Note, we did not separate via sociality given certain redundancies, e.g., the near perfect match between eusocial species and generalists. We also did not include oligolectic cavity nesting bees because there were only three species with this designation.

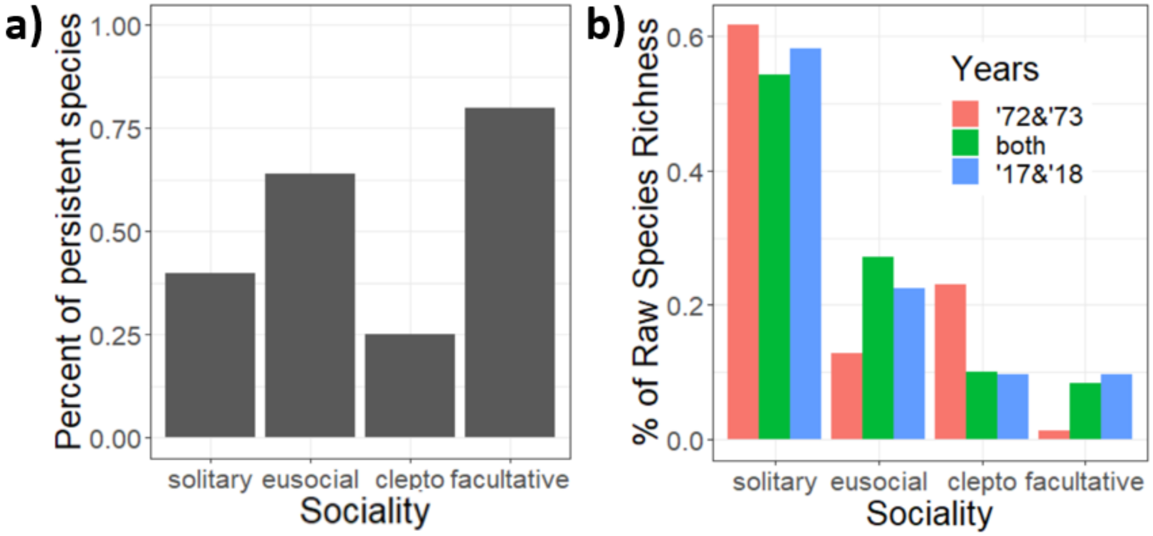


**Figure S7.** Categorical trait changes between periods of sampling bees at the E.S. George Reserve. a) Percent persistence of species separated by categorical sociality traits across raw species counts. b) Percent of raw species richness across categorical sociality traits grouped into species found only in the 70s, in both the 70s and the 2010s, or only in the 2010s. Note, sociality was not included in the neural network analysis (see Methods). Facultative bees were also not included in further analysis due to poor representation in the sample sets (~3% of historical species). Both are shown here for completeness with other categorical traits.

The neural networks trained on these categorical subsets tightened estimations of Olden importance, produced clearer connections between traits and model predictions even with low neuron counts, and revealed that initial variability seen in Fig S6 resulted from quantitative trait effects changing across categorical traits. Neural networks are typically adept at identifying such interactions without subsetting, but generally require more data than is available in our dataset. However, we show here how manual data partitioning across ecologically reasonable categories can aid in identifying patterns with smaller datasets. Results indicate that trait (predictor) effect estimates are most consistent for categories with the most extreme (high or low) chances of survival (Figs. S8, S9, S10). Oligolectic ground-nesters, polylectic cavity-nesters, and cleptoparasites all showed consistent estimates for trait effects in our networks, with oligolectic ground-nesters being the most associated with extirpation (77.8% of species extirpated), closely followed by cleptoparasites (75% extirpated). Polylectic cavity nesters were much more likely to persist (29.6% extirpated). On the other hand, 50.9% of polylectic ground nesting species persisted and had the least consistent effects' estimates (Fig. S10).

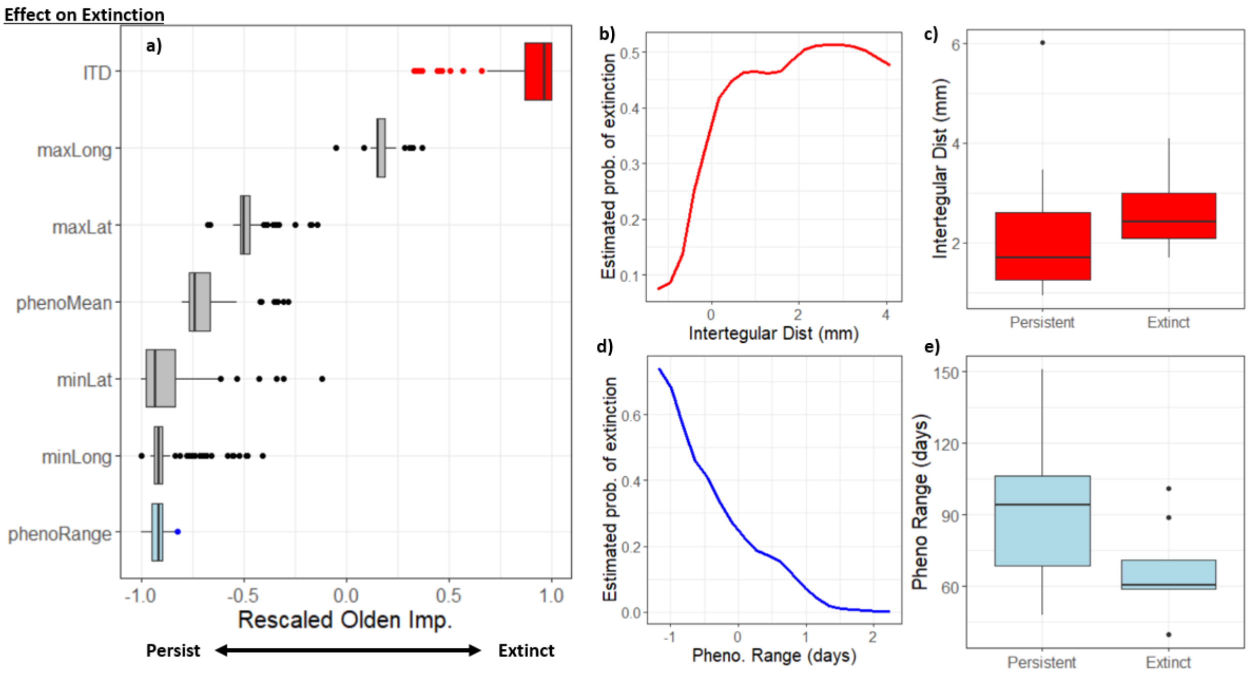


**Figure S8.** Neural network results on polylectic cavity nesting bees. a) Olden estimates of trait importance. Box and whisker plots represent 1000 iterations of neural networks trained on each categorical subset with a single hidden layer of 3 neurons and a decay rate of 5e-3. Mean accuracy was 100%. b) Example partial dependence plot showing positive relationship between ITD and extirpation probability in polylectic cavity nesters. c) Vetting ITD differences between persistent and extirpated polylectic cavity nesters. Difference is nominally significant (Kruskal Wallace test: p=0.056, $X^2$=3.65, DF=1). d) Example partial dependence plot showing positive relationship between phenological range and extirpation probability in polylectic cavity nesters. e) Vetting phenological range differences between persistent and extirpated polylectic cavity nesters. Difference is significant (t test: p=0.02, t=-2.45, DF=20.66).

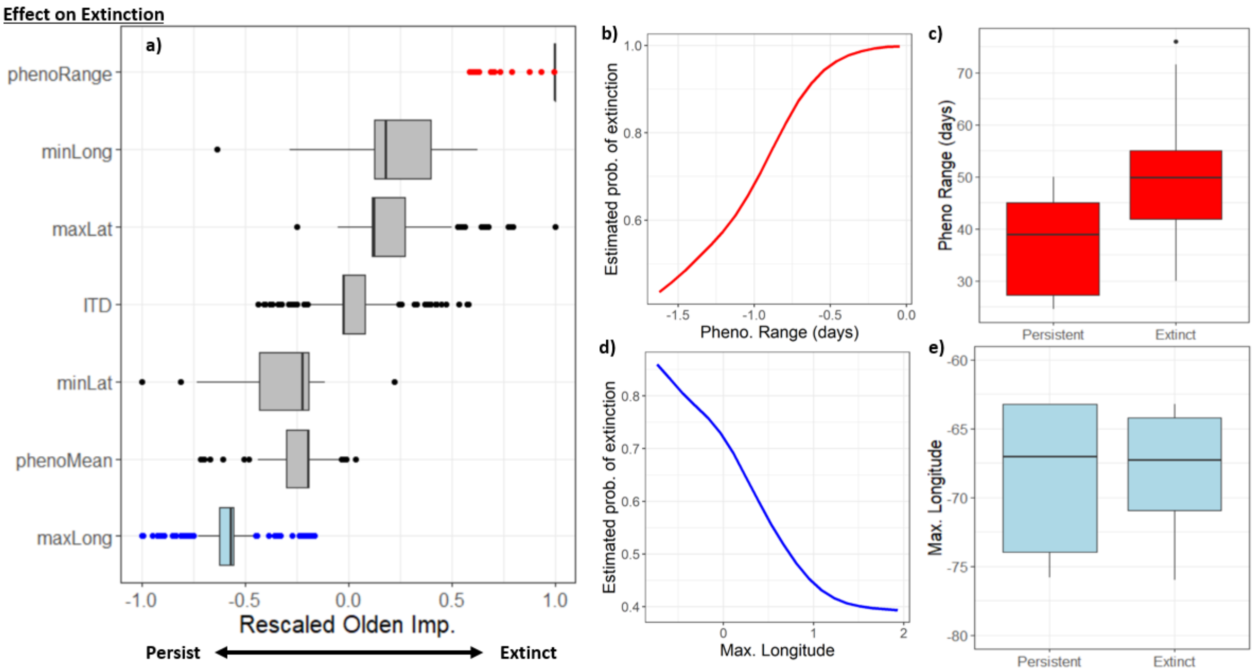


**Figure S9.** Neural network results on oligolectic ground nesting bees. a) Olden estimates of trait importance. Box and whisker plots represent 1000 iterations of neural networks trained on each categorical subset with a single hidden layer of 3 neurons and a decay rate of 5e-3. Mean accuracy was 98.8%. b) Example partial dependence plot showing positive relationship between phenological range and extirpation probability in oligolectic ground nesters. c) Vetting phenological range differences between persistent and extirpated oligolectic ground nesters. Difference is significant (t test: p=0.04, t=2.45, DF=9.07). d) Example partial dependence plot showing positive relationship between maximum longitude and extirpation probability in oligolectic ground nesters. e) Vetting maximum longitude differences between persistent and extirpated oligolectic ground nesters. Difference is not significant for all tests used.

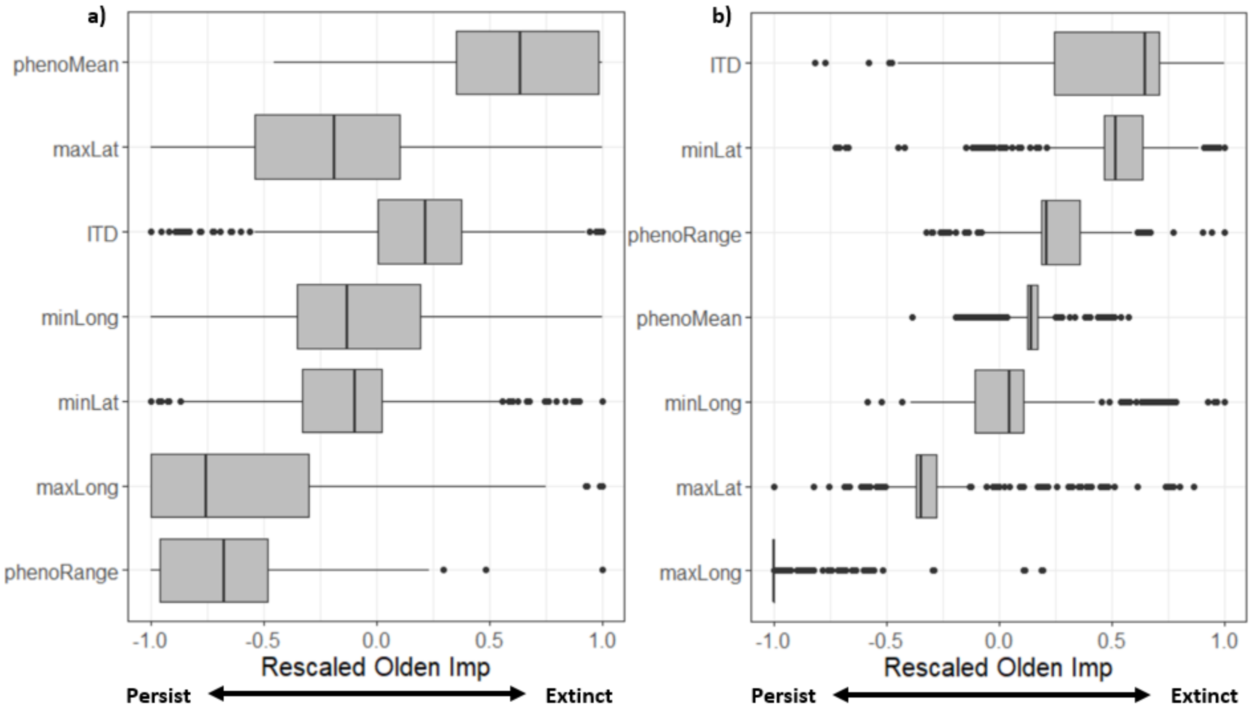


**Figure S10.** Rescaled Olden Importance values after 1000 iterations for specific diet/nesting groups of bees collected at the E.S. George Reserve. a) Polylectic ground nesters (89% accuracy across iterations) b) Cleptoparasites (mean 99.4% accuracy across iterations).

In polylectic cavity nesters (Fig. S5), the two most consistent effects were found in ITD (associated with more extirpated species) and phenological range (associated with more persistent species). For polylectic cavity nesting species, example networks produce higher extirpation probabilities as ITD increased (Fig. S8) and lower extirpation probabilities as phenological range increased (Fig. S8). Furthermore, we can “vet” our results from the traditional frequentist perspective and find that the effects hold across approaches as each of these traits are statistically different from each other between sample periods (Fig. S8). For the remaining traits, they either presented no clear monotonic effects or showed trends without statistical significance. In oligolectic ground-nesters, phenological range had a clear positive relationship with probability of extirpation (Fig. S9). This was also clear in individual neural networks and in the raw data (Fig. S9; $p=0.04$, $t=2.45$, $DF=9.07$). On the other hand, the most consistently negative relationship found between traits and extirpation probability in this category reveals the importance of "vetting" neural network results. Maximum longitude showed the strongest negative relationship, but with a mean of only ~-0.6, despite a clear trend in the partial dependence plot (Fig. S9). Investigating the relationship in our original data shows that it derives mainly from the lower minimum value in persistent species affecting the probability of extirpation in our networks (Fig. S9) rather than a significant difference between means. Also, no other trait variable presented a stronger negative relationship (Fig. S9), so by investigating the relationship in our original dataset, we can understand the limits of the relationship implied via Olden importance.

The remaining categories tested, polylectic ground-nesters and cleptoparasites, showed substantially wider ranges in effects estimates across traits (Fig S10). Despite this variability, phenological range still indicated strong effects in polylectic ground-nesters (i.e., increasing phenological range associated with increased persistence; corroborated in raw data via Kruskal-Wallis: $X^2 = 4.12$, df = 1, p = 0.04; Fig S10a). On the other hand, the cleptoparasite trait effects largely overlapped with zero and no significant relationships were discovered from any trends derived from Olden impotence values. The lack of clear monotonic relationships was at least partially driven by the higher degree of interactivity between traits (measured by H-statistic) found in both these categorical subsets (polylectic ground nesters and cleptoparasites). By investigating two-dimensional partial dependence plots, we can see examples of how trait interactions create non-linear relationships between single traits and extirpation probabilities (Fig. S11).

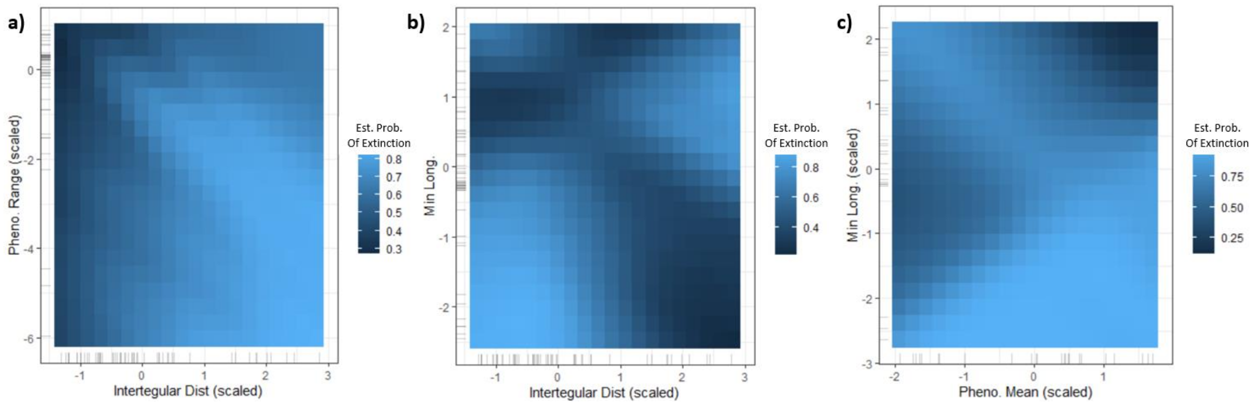


**Figure S11.** Interacting variables in neural networks used to explore trait interactions with wild bees sampled at the E.S. George Reserve. Two dimensional partial dependence plots similar to heatmaps detailing the changes in estimated probability of extinction as a function of two interacting predictor variables/network inputs. a) Interactions between ITD and phenological range in polylectic ground nesters. b) Interactions between ITD and minimum longitude in polylectic ground nesters. c) Interactions between phenological mean and minimum longitude in cleptoparasitic bees. We can see in each plot how moving along a single variable can trace increases and decreases in extinction probabilities.

At the community level, we observed clear differences between trait compositions in the historical (1972-1973) and contemporary (2017-2018) sampled communities (Fig. 2f). Two changes were made to our trait variables to make this comparison: 1) instances of each categorical trait were counted as percent of community composition, and 2) the latitude and longitude were replaced with latitudinal and longitudinal ranges to avoid negative values. Despite these changes, there are clear parallels with the neural network analysis. For example, there are significant differences along categorical variables (e.g., composition of oligolectic ground-nesting bees) and phenological range (Fig. 2f). In fact, phenological range (days of flight measured in Julian days) in the contemporary sample set is significantly larger than the historical sample for all diet/nesting categories except oligolectic, mirroring the results seen in the species based neural network analysis (Fig. S12).

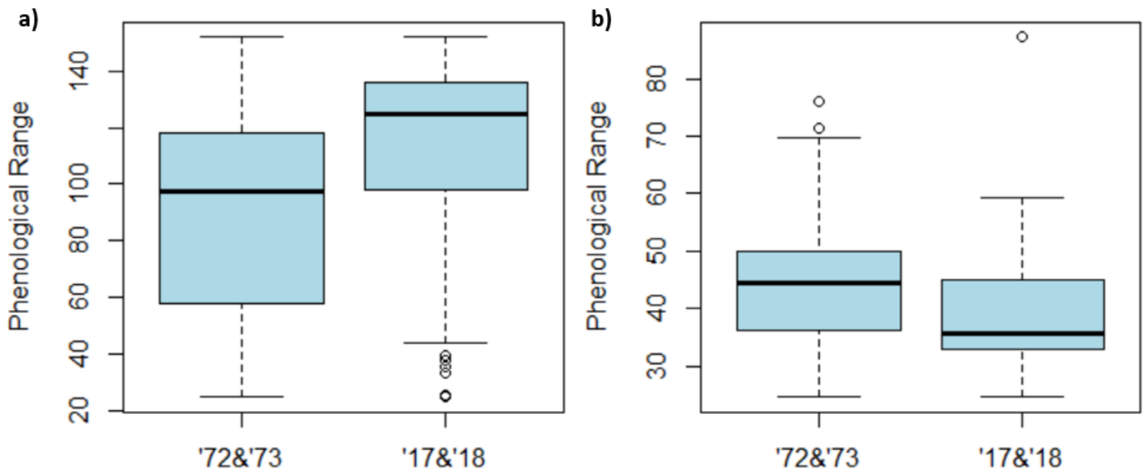

**Figure S12.** Phenological range of the bee community at the E.S. George Reserve (across sampled specimens). a) Measured across all specimens sampled in each period. Kruskal-Wallis test: chi-squared =526.98, df = 1, p < 2.2e-16. This result is consistent for all categorical subsets, b) except for oligolectic bees, in which case, the pattern is reversed (Kruskal-Wallis test: chi-squared = 20.741, df = 1, p-value = 5.257e-06).

Beyond comparisons to the neural network analysis, the results show emergent changes in community composition between periods. For example, the contemporary community of ground-nesting bees has significantly larger mean ITDs, longer mean proboscis lengths, and larger foraging distances. This trend is largely driven by the dominance of *Bombus impatiens* (Fig. S13). Interestingly, cavity nesters do not reflect the same changes, but cleptoparasitic bees do (Fig. S13). Furthermore, while we found no significant changes in latitude between periods at the species level, taking population abundance into account at the overall community level reveals significant reductions in the average maximum latitude (~3 degrees; Kruskal-Wallis $X^2$=144.84, df = 1, p<2.2e-16) and minimum latitude (~7 degrees; Kruskal-Wallis $X^2$=320.10, df = 1, p< 2.2e-16), indicating higher relative abundances of bees in the ESGR with more southerly ranges in the 2010s than the 1970s.

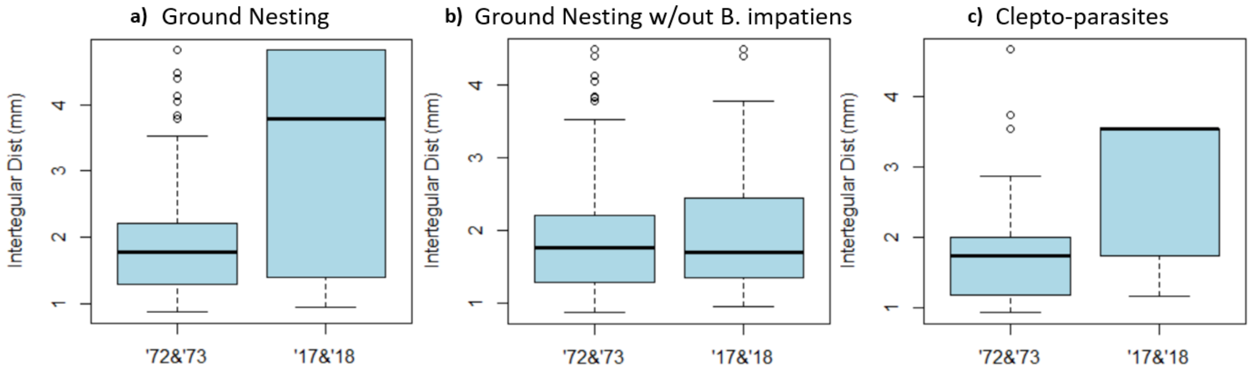


**Figure S13.** ITD changes of the bee community sampled at the E.S. George Reserve (across sampled specimens). ITD can be replaced with proboscis length or foraging distance and results would be qualitatively the same given the correlations between the three variables.

### Supplementary References


Alberdi, A. 2019. Integral Analysis of Diversity Based on Hill Numbers.

Anderson, M. J. 2001. A new method for non-parametric multivariate analysis of variance. Austral Ecology 26:32–46.

Anderson, M. J. 2004. PERMDISP: a FORTRAN computer program for permutational analysis of multivariate dispersions (for any two-factor ANOVA design) using permutation tests. Department of Statistics, University of Auckland, Auckland, New Zealand.

Bartomeus, I., J. S. Ascher, J. Gibbs, B. N. Danforth, D. L. Wagner, S. M. Hedtke, and R. Winfree. 2013. Historical changes in northeastern US bee pollinators related to shared ecological traits. Proceedings of the National Academy of Sciences of the United States of America 110:4656–60.

Basha, S. H. S., S. R. Dubey, V. Pulabaigari, and S. Mukherjee. 2020. Impact of fully connected layers on performance of convolutional neural networks for image classification. Neurocomputing 378:112–119.

Beck, M. W. 2018. NeuralNetTools: Visualization and Analysis Tools for Neural Networks. Journal of Statistical Software 85:1–20.

Bouseman, J. K., and W. E. LaBerge. 1978. A revision of the bees of the genus Andrena of the Western Hemisphere. Part IX. Subgenus Melandrena. Transactions of the American Entomological Society (1980-) 104:275–389.

Cariveau, D. P., G. K. Nayak, I. Bartomeus, J. Zientek, J. S. Ascher, J. Gibbs, and R. Winfree. 2016. The Allometry of Bee Proboscis Length and Its Uses in Ecology. PLOS ONE 11:e0151482.

Castillo, R. C. del, and D. J. Fairbairn. 2012. Macroevolutionary patterns of bumblebee body size: detecting the interplay between natural and sexual selection. Ecology and Evolution 2:46–57.

Coelho, B. W. . 2004. A review of the bee genus Augochlorella (Hymenoptera: Halictidae: Augochlorini). Systematic Entomology 29:282–323.

Evans, F. C. 1986. Bee - Flower Interactions on an Old Field in Southeastern Michigan. Pages 103–109 *in* G. K. Clambey and R. H. Pembley, editors. The prairie: past, present and future. Proceedings of the ninth North American Prairie Conference. Tri-College University Center for Environmental Studies, Fargo, ND.

Gibbs, J. 2011. Revision of the metallic Lasioglossum (Dialictus) of eastern North America (Hymenoptera: Halictidae: Halictini). Zootaxa 3073:1–216.

Gibbs, J., E. Hanuschuk, R. Miller, M. Dubois, M. Martini, S. Robinson, P. Nakagawa, C. Sheffield, S. Cardinal, and T. Onuferko. (n.d.). A checklist of the bees (Hymenoptera: Apoidea) of Manitoba. The Canadian Entomologist.

Gibbs, J., L. Packer, S. Dumesh, and B. N. Danforth. 2013. Revision and reclassification of Lasioglossum (Evylaeus), L. (Hemihalictus) and L. (Sphecodogastra) in eastern North America (Hymenoptera: Apoidea: Halictidae). Zootaxa 3672:1–117.

Gorman, B. 2018. mltools: Machine Learning Tools.

Greenleaf, S. S., N. M. Williams, R. Winfree, and C. Kremen. 2007. Bee foraging ranges and their relationship to body size. Oecologia 153:589–596.

Heaton, J. 2008. Introduction to neural networks with Java. Heaton Research, Inc.

Hung, K.-L. J. 2017. Effects of Habitat Fragmentation and Introduced Species on the Structure and Function of Plant-Pollinator Interactions. University of California, San Diego, San Diego, CA.

IUCN Standards and Petitions Subcommittee. 2014. Guidelines for Using the IUCN Red List Categories and Criteria. IUCN.

Kendall, L. K., R. Rader, V. Gagic, D. P. Cariveau, M. Albrecht, K. C. R. Baldock, B. M. Freitas, M. Hall, A. Holzschuh, F. P. Molina, J. M. Morten, J. S. Pereira, Z. M. Portman, S. P. M. Roberts, J. Rodriguez, L. Russo, L. Sutter, N. J. Vereecken, and I. Bartomeus. 2019. Pollinator size and its consequences: Robust estimates of body size in pollinating insects. Ecology and Evolution 9:1702–1714.

Kuhn, M. 2015. caret: Classification and Regression Training.

LaBerge, W. E. 1971. A revision of the bees of the genus Andrena of the Western Hemisphere. Part IV. Scrapteropsis, Xiphandrena and Raphandrena. Transactions of the American Entomological Society 97:441–520.

LaBerge, W. E. 1973. A revision of the bees of the genus Andrena of the Western Hemisphere. Part VI. Subgenus Trachandrena. Transactions of the American Entomological Society 99:235–371.

LaBerge, W. E. 1977. A revision of the bees of the genus Andrena of the Western Hemisphere. Part VIII. Subgenera Thysandrena, Dasyandrena, Psammandrena, Rhacandrena, Euandrena, Oxyandrena. Transactions of the American Entomological Society 103:1–143.

LaBerge, W. E. 1980. A revision of the bees of the genus Andrena of the western hemisphere. Part X. Subgenus Andrena. Transactions of the American Entomological Society 106:395–525.

LaBerge, W. E. 1985. A revision of the bees of the genus Andrena of the Western Hemisphere. Part XI. Minor subgenera and subgeneric key. Transactions of the American Entomological Society 111:441–567.

LaBerge, W. E. 1989. A revision of the bees of the genus Andrena of the Western Hemisphere. Part XIII. Subgenera Simandrena and Taeniandrena. Transactions of the American Entomological Society 115:1–56.

LaBerge, W. E., and J. K. Bouseman. 1970. A revision of the bees of the genus Andrena of the Western Hemisphere. Part III. Tylandrena. Transactions of the American Entomological Society 4:543–605.

Lerman, S. B., and J. Milam. 2016. Bee Fauna and Floral Abundance Within Lawn-Dominated Suburban Yards in Springfield, MA. Annals of the Entomological Society of America

109:713–723.
Lim, K., S. Lee, M. Orr, and S. Lee. 2022. Harrison's rule corroborated for the body size of cleptoparasitic cuckoo bees (Hymenoptera: Apidae: Nomadinae) and their hosts. Scientific Reports 2022 12:1 12:1–12.
McGinley, R. J. 1986. Studies of Halictinae (Apoidea: Halictidae), I: Revision of New World Lasioglossum Curtis. Smithsonian Contributions to Zoology 429:1–294.
Mitchell, T. B. 1960. Bees of the Eastern United States: volume I. N. C. Agric. Exp. Stn. Tech. Bull. 141:1–538.
Mitchell, T. B. 1962. Bees of the Eastern United States: volume II. N. C. Agric. Exp. Stn. Tech. Bull. 152:1–557.
Molnar, C. 2022. iml: Interpretable Machine Learning.
Nicholson, C. 2019. Nicholson_etal_2019_Wild_bee_occurrence_traits. figshare.
Normandin, É., N. J. Vereecken, C. M. Buddle, and V. Fournier. 2017. Taxonomic and functional trait diversity of wild bees in different urban settings. PeerJ 2017:e3051.
Oksanen, J., F. G. Blanchet, M. Friendly, R. Kindt, P. Legendre, D. McGlinn, P. R. Minchin, R. B. O'Hara, G. L. Simpson, P. Solymos, M. H. H. Stevens, E. Szoecs, and H. Wagner. 2019. vegan: Community Ecology Package.
Rehan, S. M., and C. S. Sheffield. 2011. Morphological and molecular delineation of a new species in the Ceratina dupla species-group (Hymenoptera: Apidae: Xylocopinae) of eastern North America 1. Zootaxa 2873:35–50.
Ribble, D. W. 1967. The Monotypic North American Subgenus Larandrena of Andrena (Hymenoptera: Apoidea). Bulletin of the University of Nebraska State Museum 6:27–42.
Salinas, H. 2021. ecolTest: Community Ecology Tests.
Venables, W. N., and B. D. Ripley. 2002. Modern Applied Statistics with S. Fourth. Springer, New York.
Wickham, H., and D. Seidel. 2022. scales: Scale Functions for Visualization.
Wood, S. 2022. mgcv: Mixed GAM Computation Vehicle with Automatic Smoothness Estimation.
Wood, S. N. 2004. Stable and efficient multiple smoothing parameter estimation for generalized additive models. Journal of the American Statistical Association 99:673–686.
Wood, S. N. 2011. Fast stable restricted maximum likelihood and marginal likelihood estimation of semiparametric generalized linear models. Journal of the Royal Statistical Society: Series B (Statistical Methodology) 73:3–36.
